# Predictions of spatio-temporal patterns of soil respiration in a mixed forest: Downscaled meteorological data outperform local parameters

**DOI:** 10.64898/2026.01.21.700858

**Authors:** Julian Brzozon, Friederike Lang, Patricia Schwarzkopf, Teja Kattenborn, Julian Frey, Helmer Schack-Kirchner

**Affiliations:** Chair of Soil Ecology, University of Freiburg, Bertoldstrasse 17, 79098 Freiburg, Germany; Chair of Sensor-based Geoinformatics, University of Freiburg, Tennenbacher Str. 4, 79106 Freiburg, Germany; Chair of Forest Growth and Dendroecology, University of Freiburg, Tennenbacher Str. 4, 79106 Freiburg, Germany

**Keywords:** Tree distribution, atmospheric vapour pressure, time-lag effects, hot spots, hot moments

## Abstract

**Introduction:** Patterns of soil respiration (R_s_) are heterogeneous on spatio-temporal scales. The best studied controlling factors of R_s_ are microclimatic conditions such as soil temperature and moisture. However, a strong pronounced seasonality shifts R_s_ patterns from temperature to moisture-controlled regimes. Hereby, lag effects and influences of trees in mixed forests on large spatio-temporal scales are rarely investigated.

**Material and Methods:** We investigated Rs over two years on a weekly to fortnightly measurement frequency at a 1 ha area in a mixed forest on 35 predefined locations using open bottom chambers. Analysis was derived using downscaled meteorological data and a tree species map.

**Results:** Our data confirmed soil temperature and moisture as important variables which explained up to 72% in variability. Yet, our results revealed an additional predictor: the atmospheric vapour pressure. Together with seasonality and the vapour pressure deficit, 84% in variability could be explained. Additionally, the predictors traced a decrease in Rs due to drought and an increase due to rewetting. By tendency, Rs correlated negatively with distance to the tree and Rs was significantly higher in broadleaf patches compared to coniferous and mixed patches during the summer seasons.

**Conclusion:** We conclude that meteorological conditions might serve as valuable predictors for CO2 emissions from forest soil. This enables upscaling of Rs given the dense availability of meteorological data. Tree species distribution partly explained the spatial patterns of Rs. However, a detailed analysis of local soil properties would enhance our understanding of the interactions between soils, plants and the atmosphere.

## 1. Introduction

Soil respiration (R_s_), the release of carbon dioxide (CO_2_) from soils, represents the second largest terrestrial CO_2_ flux and the biggest contributor to total ecosystem respiration (Goulden et al., 1996; Longdoz et al., 2000). Even though temporal patterns showed similar trends, independent measurements of R_s_ yielded higher fluxes compared to ecosystem respiration at various ecosystem scales (Barba et al., 2018). A certain proportion of this discrepancy may be explained by heterogeneity of R_s_ in ecosystems, as well as the existence and abundance of hotspots (Barba et al., 2018; Hereș et al., 2021; Scapucci et al., 2025). R_s_ is composed of heterotrophic respiration (R_h_) and autotrophic respiration (R_a_) (Kuzyakov, 2006), while R_a_ is seen as the main contributor to total R_s_ (Raich and Tufekciogul, 2000). Thus, the big spatio-temporal variability of R_s_ may be controlled by the ecosystem structure (Lejon et al., 2005; Søe and Buchmann, 2005; Štursová et al., 2016).

Across European forest ecosystems, only few, mostly not current studies, examined specifically spatio-temporal patterns of R_s_ at broader scales using the standard open bottom chamber technique (Table S1). Comparisons between the spatial scales are difficult due to a lack of standardization in study designs (number of plots and measurements per plot). Furthermore, available research depicts the trade off between temporal and spatial scale in R_s_ measurements. Large-scaled spatial measurement campaigns are usually spread out over several days as it is hardly feasible to complete more than a certain number of measurements within one day. Automated chambers offer the possibility to measure on a high temporal scale but usually cover only single point measurements. Furthermore, permanent chambers on the soil may bias R_s_ rates (Ruehr et al., 2010). Nevertheless, literature summarised known controlling variables for R_s_, which are mostly microclimatic conditions such as soil temperature (T_s_) or soil water content (SWC) (Hereș et al., 2021; Ruehr et al., 2010; Schindlbacher et al., 2012). A widely accepted relationship between R_s_ and T_s_ can be described by site-specifically fitted exponential functions, which can often explain high proportions of variability in R_s_. From these functions, the increase in R_s_ by changes in T_s_ of 10 °C can be derived (Q_10_ value). However, a simple use of this value neglects seasonality (Janssens and Pilegaard, 2003) and important interaction effects with SWC (Lloyd and Taylor, 1994; Qi et al., 2002). Even though R_s_ models based on T_s_ perform well, a rather minor (Ekblad et al., 2005) or indirect (Kuzyakov and Gavrichkova, 2010) role of T_s_ was suggested due to time lag-effects. An increase in solar radiation during spring, peaking during the summer season (in temperate ecosystems), also drives changes in T_s_ (Kuzyakov and Gavrichkova, 2010). Such meteorological conditions prior to R_s_ measurements represent rarely investigated control variables for R_s,_ but they might play a crucial role, as they drive ecosystem activity. This highlights the importance of plants as contributors to R_s_ in ecosystems due to the dependency of R_a_ on photosynthesis (Kuzyakov and Gavrichkova, 2010; Moyano et al., 2008; Ruehr and Buchmann, 2010; Tang et al., 2005a). Ekblad et al. (2005) and Bowling et al. (2002) highlighted the importance of lag-effects of the vapour pressure deficit (VPD). These are, to the current state of knowledge, the only studies using the VPD to explain patterns of isotopic signatures (^13^C) of R_s_, even though the VPD might be used as a proxy for ecosystem stress due to plant-specific stomatal behaviour as a response to changes in VPD (e.g. Hsiao, 1973; McAdam and Brodribb, 2015; Sanginés de Cárcer et al., 2018). However, at given air temperature (T_a_), relative air humidity (H_rel_) is the determining factor for the VPD. H_rel_ is rarely mentioned in the literature but may affect the physiological status of trees and understorey vegetation in a forest ecosystem (Kupper et al., 2011). Kukumägi et al. (2014) presented reductions in R_s_ due to almost permanently increased H_rel_ in a hemiboreal forest ecosystem. However, shorter-term lag effects of 2-4 hours were also observed for microclimatic parameters such as T_s_, depending on soil depth (Ruehr et al., 2010). However, Mitra et al. (2019) reported diurnal dependence of R_s_ on photosynthetic active radiation (1.5-3 hours) while temperature and SWC determined variability on intermediate (8-15 days) up to longer (15-30 days) time scales. Therefore, available research implies a multiple, complex dependency of R_s_ on microclimatic but also on meteorological conditions.

On spatial scale, forest structure can be of crucial control for patterns in R_S_. In a mixed forest with a high density of various tree species, Vesterdal et al. (2012) observed the highest R_s_ associated with European beech trees. In a mixed forest containing both, pure patches and mixed patches, Khomik et al. (2006) observed the highest R_s_ associated with deciduous compared to coniferous patches, as well as a higher R_s_ associated with deciduous trees within the mixed patches. Similar to these results, Longdoz et al. (2000) reported higher R_s_ associated with European beech compared to Douglas fir patches within one forest. Jochheim et al. (2022) also report differences between deciduous and coniferous patches, but these varied between the measurement years depending on drought conditions. In an early literature review, Raich and Tufekciogul (2000) summarized in general 10 % lower R_s_ in coniferous forests compared to deciduous forests. These contrasts in R_s_ associated with species composition may be due to tree specific habitats, as associated root systems (Köstler et al., 1968), microbial and fungal community compositions (e.g. Hofmann et al., 2023; Prescott and Grayston, 2013) and nutrient availability (e.g. Spielvogel et al., 2016; Vesterdal, 1999; Vesterdal et al., 2013) depend on the tree species. Additionally, seasonal differences in photosynthetic activity between evergreen coniferous trees and the seasonal change in foliation of deciduous trees (Hansen et al., 1997; Jochheim et al., 2022; Longdoz et al., 2000) are important drivers for spatio-temporal patterns of R_s_. While several studies address tree species effects, only few of them highlight the effects of distance to the stemfoot on R_s_ as controlling variable. Søe and Buchmann (2005) discovered no significant relationship between the distance to the closest tree and R_s_ rate in a European beech forest. However, Jochheim et al. (2022) observed constantly the highest R_s_ closest to the trees (1.25 m), compared to more distant positions (2.5 and 3.5 m), with more pronounced species-specific effects during the summertime compared to the wintertime in pure stocks. In a higher elevated subalpine coniferous forest, Scott-Denton (2003) determined a negative correlation between the distance to the tree and the associated R_s,_ which was, however, more likely to be explained by the forest floor thickness. In a Sitka pine stand, Saiz et al. (2006b) reported higher R_s_ and stronger fluctuations at positions closer to the tree. Similar results were reported for Loblolly pine stands, although not at this fine spatial resolution (Wiseman and Seiler, 2004). A further positive correlation regarding forest structural features was determined for the average diameter at breast height (DBH) of trees in proximity to the measurement position (Søe and Buchmann, 2005).

Due to these complex dependencies of R_s_ on spatio-temporal patterns, future trends of total R_s_ in the scope of climate change with extreme weather conditions such as heavy rain or extended drought periods are ambiguous (IPCC, 2023). Especially dry periods can alter the pattern of R_s_ with collapses during drought and increases due to rewetting, which was early discovered as the “Birch-effect” (Birch, 1958). However, the effects of changes in meteorological conditions on R_s_ are uncertain, because contributors to R_s_ as microorganism activity (Schindlbacher et al., 2011; Schindlbacher et al., 2012), root distribution (Brunn et al., 2022; Brunner et al., 2015) as well as spatially variable substrate availability (Brunn et al., 2022; Brunn et al., 2023), might alter with climate change. The goal of this study is to obtain reliable data for R_s_ from a mixed forest, to determine its controlling factors and to quantify the contribution of predictor variables on R_s_. Based on the state of knowledge we hypothesize that:

1. temporal patterns of R_s_ depend on T_s_ but an additional proportion in variation can be explained by more or less lagged meteorological parameters as T_a_, atmospheric moisture conditions (VPD, VP_cur_, H_rel_) and global radiation.
2. Hot moments such as a pulse (“Birch effect”) after rewetting may contribute significantly to total R_S_.
3. Tree distribution causes spatio-temporal variability. With tree specific phenology, R_s_ associated with deciduous trees is higher during the summer season, while R_s_ associated with coniferous trees is higher during the winter season. We expect higher R_s_ closer to tree stems compared to R_s_ with a larger distance to tree stems, due to respiratory processes of trees.

To test our stated hypothesises, we measured R_s_ on an area of around 10000 m^2^ at 35 predefined positions over 2 years and fitted those into a statistical model. This approach had also the aim to identify hot spots and hot moments in R_s_. Following, hot spots (Stutz and Lang, 2023) were defined as locations that, on average across the entire area, reflect significantly higher R_s_ while cold spots are locations with significantly lower R_s_ than the seasonal averages. Hot moments were defined phases in the temporal development of R_s_, that deviate from the pattern with unusual reactions (McClain et al., 2003), which were defined as changes in known controlling variables (T_S_). Our investigations are part of the collaborative research center ECOSENSE (Werner et al., 2024) with its joint research area (ECOSENSE Forest).

## 2. Material & Methods

### 2.1. Research area

The ECOSENSE Forest is located in the south-western foothills of the Black Forest in Germany (48.2684, 7.8779, WGS84) on an average elevation of 465-475 m a.s.l.. Dominant tree species are European beech (*Fagus sylvatica* L.) and Douglas fir (*Pseudotsuga menziesii* Mirbel). Silver fir (*Abies alba* Mill.), European larch (*Larix decidua* Mill.) and Norway spruce (*Picea abies* L.) occur sporadically, while understorey vegetation is only sparsely developed. The entire stock is approximately 55-110 years old. The long-year (1981 – 2010) mean precipitation derived from the HYRAS dataset (compare chapter 2.5.) is 1000 mm yr^-1^, with a mean annual temperature of 10°C (DWD, 2025).

### 2.2. Soil Properties

Full description of soil is based on two pits excavated in November 2024 according to IUSS (2022). Bulk samples were taken from the corresponding depth directly next to undisturbed rings (200 cm^3^). Due to the high skeleton content in the northern profile, it was not possible to extract any undisturbed rings below a depth of 42 cm and physical parameters were therefore not determinable (n.d.).

The organic forest floor consisted of a patchy f-mull type with an averaging thickness of 2-4 cm. Two geological parent materials were present, resulting in two different soil types. In the southern part of the research area, we identified a Dystric Stagnic Cambisol (loamic) (IUSS, 2022) derived from carbonate-free quaternary loess covering mesozoic sediments that are strongly weathered. Redoximorphic features occur below a depth of 40 cm. In the northern part of the research area, we identified a Dystric Skeletic Cambisol (siltic, humic) (IUSS, 2022) derived from carbonate-free quaternary loess, covering mesozoic sandstone with a high skeleton content. Below a depth of 40 cm only thin layers of fine earth between platy stones can be found. Soil was analysed according to standard methods (Table S2) for physical and chemical properties (Table 1).

**Table 1:** Soil chemical and physical properties. Values represent mean values ± standard deviation.

| Soil type | depth [cm] | pH<sub>H2O</sub> [-] | C<sub>org</sub> [mg g<sup>-1</sup>] | C:N [-] | CEC [μmol<sub>c</sub> g<sup>-1</sup>] | D<sub>s</sub>/D<sub>0</sub> [-] | $\phi_{soil}$ [-] | $\rho_{bulk}$ [g cm<sup>-3</sup>] | CM [%vol] | Clay [%] | Particle size | |
| --- | --- | --- | --- | --- | --- | --- | --- | --- | --- | --- | --- | --- |
|  |  |  |  |  |  |  |  |  |  |  | Silt [%] | Sand [%] |
| Dystric | 0-7 | 4.8 | 21.6 ± 0.9 | 11.8 ± 0.2 | 74 ± 2 | 0.07 ± 0.02 | 0.56 ± 0.05 | 1.15 ± 0.07 | 0 | 34 ± 1.5 | 59 ± 1.2 | 7 ± 0.3 |
| Stagnic | 7-25 | 4.5 | 9.5 ± 0.1 | 9.3 ± 0.2 | 73 ± 3 | 0.05 ± 0.02 | 0.53 ± 0.05 | 1.24 ± 0.09 | 0 | 33 ± 0.6 | 60 ± 0.4 | 7 ± 0.2 |
| Cambisol (loamic) | 25-38 | 4.6 | 3.6 ± 0.1 | 5.1 ± 0.1 | 96 ± 1 | 0.05 ± 0.02 | 0.51 ± 0.04 | 1.37 ± 0.1 | 3 | 39 ± 0.4 | 54 ± 0.8 | 7 ± 0.4 |
| | 38-60 | 4.8 | 3.1 ± 0.4 | 4.7 ± 0.4 | 104 ± 3 | 0.05 ± 0.01 | 0.45 ± 0.04 | 1.54 ± 0.1 | 3 | 42 ± 0.4 | 52 ± 0.1 | 6 ± 0.5 |
| | 60-80 | 4.8 | 1.5 ± 0.1 | 2.4 ± 0.3 | 100 ± 1 | 0.01 ± <0.01 | 0.38 ± 0.02 | 1.69 ± 0.07 | 8 | 44 ± 0.5 | 53 ± 0.7 | 4 ± 0.2 |
| | 80-100 | 4.9 | 0.5 ± <0.1 | 0.9 ± 0.1 | 87 ± 1 | 0.01 ± <0.01 | 0.35 ± 0.01 | 1.79 ± 0.04 | 3 | 35 ± 0.5 | 58 ± 0.6 | 7 ± 0.3 |
| Dystric | 0-7 | 4.3 | 25.7 ± 0.6 | 14.5 ± 0.5 | 75 ± 2 | 0.06 ± 0.02 | 0.57 ± 0.04 | 1.07 ± 0.09 | 0 | 26 ± 0.2 | 57 ± 0.4 | 17 ± 0.3 |
| Skeletic | 7-20 | 4.4 | 13.3 ± 0.4 | 12.0 ± 0.7 | 59 ± 3 | 0.07 ± 0.02 | 0.54 ± 0.03 | 1.22 ± 0.08 | 0 | 25 ± 0.3 | 58 ± 0.5 | 18 ± 0.3 |
| Cambisol (siltic, humic) | 20-42 | 4.4 | 11.0 ± 1.7 | 12.0 ± 1.4 | 56 ± 2 | 0.07 ± 0.03 | 0.53 ± 0.05 | 1.24 ± 0.1 | 17 | 27 ± 0.9 | 54 ± 0.8 | 19 ± 0.5 |
| | 42-100 | 4.3 | 4.7 ± 0.4 | 6.7 ± 0.5 | 75 ± 2 | n.d. | n.d. | n.d. | 63 | 35 ± 0.5 | 49 ± 0.5 | 16 ± 0.3 |

Abbreviations: organic carbon (C<sub>org</sub>), carbon to nitrogen ratio (C:N), effective cation exchange capacity (CEC), relative diffusion coefficient (D<sub>s</sub>/D<sub>0</sub>) at -300 hPa water potential, porosity ( $\phi_{soil}$ ), bulk density ( $\rho_{bulk}$ ) and coarse material (CM)

### 2.3. Soil respiration measurements

R_s_ was measured manually over two years on an approximately weekly-to bi-weekly base (January 2023-December 2024) on 35 plots using the open bottom chamber technique. Each of the measuring plots as basic spatial investigation unit had an area of 2.25 m^2^. The plots were established with distances of 10 m between each plot on a north-south axis (21 plots) flanked by 20 m off-transect points east and west (14 plots). The plots covered the approximate main area of the ECOSENSE Forest and are on a ridge with little variability in terrain slope. For each measurement two open bottom chambers (weight: ca. 1680 g, volume: 0.0018 m^3^, soil-coverage area: 0.022 m^2^) each equipped with a solid-state CO_2_-probe (Vaisala GMP343, Vaisala Oyj, Helsinki, Finland) were placed directly on the forest floor at a random position within the plot. The individual measurements on every plot were averaged to one measurement and reflect therefore the R_s_ on plot level. After each measurement, the chambers were left open exposed to the atmosphere to reach near-ambient conditions (Fiedler et al., 2022). To avoid any bias caused by diurnal patterns, the order of measurement plots was determined at random each day (Ruehr et al., 2010). Chambers were placed on the floor without collars (Snell et al., 2014) to avoid artifacts and disturbance due to the invasive procedure of collar insertion (Jovani-Sancho et al., 2017; Wang et al., 2005). To compensate for lateral leakage, a correction factor was calculated based on time series with Neon injection as tracer gas on a simulated forest floor surface in the lab (Equation 1).

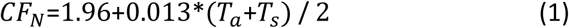

With *T_a_* = air temperature in °C and *T_s_* = soil temperature in °C.

From June 2023 we additionally placed a combination of a chain and a rubber-foam sealing (weight: ca. 1750 g) between the base of the chambers and soil surface and calculated a separate correction factor (Equation 2).

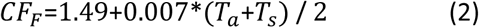

The measurement time was adapted to the expected flux (Fiedler et al., 2022): at the beginning of the measurement campaign, we started measurements with a length of 390 seconds (20 measurements, interval of 20 seconds). Initial analysis revealed that closing time could be reduced and was set to 190 seconds (20 measurements, gap of 10 seconds). Since the measurement time was shortened, we were able to measure all plots within one day (since June 2023). For R_s_ flux calculation 50 seconds were cut off at the beginning and at the end of each single measurement due to saturation effects at the end or slow increase at the beginning. On three days, measurements were not completed due to weather conditions or system error (once only 33/35 plots and twice only 26/35 plots were measured). Due to technical problems in 4% out of all measurements, R_s_ was measured only with one chamber. During R_s_-measurements, T_s_ was measured in ca. 13 cm soil depth (GMH 3700 GES401, Seseca GmbH, Germany) and volumetric SWC (SM150T, Delta-T Devices, UK) in ca. 5 cm depth. Unfortunately, the measurement device for SWC broke down in January 2024 due to high skeleton content and measurements were not possible thereafter.

### 2.4. Forest stock features

A forest inventory was conducted in the core area of the ECOSENSE Forest (ca. 3 ha; Werner et al., 2024). All trees with a diameter at breast height (DBH) ≥ 7 cm were recorded in the field, including species identity, DBH, and spatial position. Tree locations were initially determined using an EMLID RTK GNSS system (EMLID REACH RS2, Emlid Tech Kft. Budapest, Hungary). Subsequently, terrestrial laser scanning (TLS) was carried out using a Riegl VZ-400i (RIEGL Laser Measurement Systems GmbH, Horn, Austria) along multiple transects to capture the stand structure in high detail (3Dtrees.earth, 2025). The resulting point clouds were used to refine (optimize) the geolocation of individual tree positions and to derive tree heights for the complete inventory. The R_s_ measurement plots were triangulated to a random position within the plot to the nearest neighbouring trees. Additionally, we buffered a zone around the measurement position (5 m radius) and calculated the average distance to all trees, the number of trees and the average DBH within the buffered area. We used the distance of 5 m to include all plots into analysis, as the maximum nearest neighbouring distance was 4.5 m. As measurements for R_s_ were performed randomly on a 2.25 m^2^ area, the exact position including distance to trees has a precision range of ± 1 m. Therefore, the distance to the nearest tree was classified into four categories each covering 1.25 m with increasing distance. This covers also the selected distances represented in literature (e.g. Jochheim et al., 2022). Since not only *Fagus sylvatica* and *Pseudotsuga menziesii* were within the buffer zone we grouped the tree species according to the species composition, which are also referred to as these groups in the following: deciduous (pure *Fagus sylvatica* composition [18 plots]), coniferous (either pure *Pseudotsuga menziesii* [8 plots], pure *Abies alba* [1 plot] or *Pseudotsuga menziesii* mixed with *Abies alba* [2 plots]) and mixed (always containing *Fagus sylvatica* mixed with different contributions of *Pseudotsuga menziesii, Abies alba, Larix decidua, Picea abies* [6 plots]). The number of trees within the buffer zone varied between 1-10 trees.

### 2.5. Meteorological data

We downloaded freely available meteorological data (HYRAS v6-1) from the German weather service (DWD, 2025) which represents modelled data of daily precipitation (Rauthe et al., 2013), minimum, maximum and average air temperature (T_a_) and relative humidity (H_rel_) (Frick et al., 2014; Razafimaharo et al., 2020), downscaled for the specified cell covering the ECOSENSE Forest (5x5 km cell-size). The data are historically available from 1931 until present. We calculated the saturated vapour pressure (VP_sat_) using Teten’s equation. The current vapour pressure (VP_cur_) was calculated by multiplying VP_sat_ by H_rel_. The atmospheric vapour pressure deficit (VPD) was calculated by subtraction of VP_cur_ from VP_sat_. Wind velocity and global radiation data were extracted from the ERA5-Land dataset (Copernicus Climate Change Service, 2019; Muñoz-Sabater et al., 2021), which also represents a modelled grid (11 x 7 km cell size). As a smooth model parameter of seasonality we calculated the negative cosine of the Julian day of the year (cos_doy_) (e.g. Sims, 1974), with positive, increasing values during the mid of the year (+1) and decreasing, negative values during the start and end of the year (-1). We also calculated the negative sine of the Julian day of the year (sin_doy_), which then creates the same effect but shifted by three months, and therefore represents the spring and autumn season. This method takes into account a transition with a development, apart from standard seasonal classification, which cumulates three months to only one variable. In order to obtain more general statements about the seasonal effects we also used the seasonal classification according to the German weather service (DWD) and averaged the values over the seasons for a more common evaluation. Seasons were classified as Winter: December – February, Spring: March – May, Summer: June – August, Autumn: September – November.

### 2.6. Statistical analysis

To determine the predictors for **temporal patterns** in R_s_ we used linear models (*lm* from stats-package, R Core Team, 2023) applied to the daily R_s_ as average from all 35 sampling plots. As predictors, we used the respective average meteorological parameters from the day of the measurements of R_s_ as well as aggregated mean or summarised values of these predictors 1-7 days prior to the measurement day. To prevent model overfitting and to identify predictors that provide meaningful information on the development of R_s_, only predictors with an R^2^ above 0.4 were taken into consideration for the subsequent model optimization. Additionally, we also applied three common approaches to determine temporal patterns by checking dependencies of R_s_ on T_s_ alone (Equation 3) and in combination with SWC (Equation 4). We also calculated Q_10_ values (Equation 5) using non least square models (*nls* from stats-package, R Core Team, 2023) with fitted parameters.

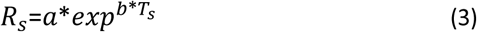

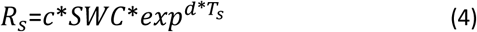

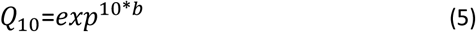

With *R_s_* = the modelled soil respiration [µmol CO_2_ m^-2^ s^-1^], *T_s_* = soil temperature [°C] at reference depth (13 cm) and *SWC* [%_vol_]. *a*, *b*, *c*, *d* were fitted using non least square models.

We used variance inflation factor (VIF) to rule out collinearity among the predictors (*vif* from car, Fox and Weisberg, 2019) and used only combinations in which the VIF was < 3 for all predictors (O’brien, 2007). Model residuals were tested for autocorrelation using the Breusch-Godfrey test (Godfrey, 1978) (*bgtest* from lmtest-package Achim Zeileis and Torsten Hothorn, 2002). For the nls models, local linearization was used prior to the application of the Breusch-Godfrey test. In case of of heteroscedasticity of residuals, which were tested using diagnostic plots, robust standard errors (HC3) were calculated for linear models (*vcov* from stats-package R Core Team, 2023) and for nls models (*glns* from nlme-package Pinheiro and Bates, 1999). To visualize and determine **spatial patterns,** we used inverse distance weighting (*idw* from phylin-package, Tarroso et al., 2019). Dependencies of R_s_ on forest structural features were tested using linear models (*lm* from from stats-package, R Core Team, 2023), spearman correlation (*cor.test* from stats-package, R Core Team, 2023) and pairwise wilcox rank sum tests (from stats-package, R Core Team, 2023) for group comparisons. Answering the question whether (seasonal) **hot or cold spots** exist in the ECOSENSE Forest, we calculated a mixed effects model (*lmer* from lme4, Bates et al., 2015) with the measurement plot as explanatory variable and the day of the measurement as a random effect. From this, we calculated estimated marginal mean values (*emmeans* from emmeans package, Lenth, 2023) and set those in contrast (*contrast* from stats-package, R Core Team, 2023) to the averaged values of the corresponding season, to identify plots with statistically significant higher or lower deviations from the averaged seasonal R_s_. To identify **hot moments**, we used linear models (*lm* from stats-package, R Core Team, 2023) and spearman correlation tests (*cor.test* from stats-package, R Core Team, 2023) to check for unusual patterns between R_s_ and T_s_. For data organization we used the packages tidyverse (Wickham et al., 2019) and lubridate (Grolemund and Wickham, 2011), for visualization ggplot2 (Wickham, 2016). All statistical analysis was done in RStudio 2023.06.0 based on R4.2.3 (R Core Team, 2023).

## 3. Results

### 3.1. Weekly to fortnightly soil respiration measurements

In total, 4507 single R_s_ measurements were available for data evaluation, which were averaged to 2295 measurements on the plot level on a total of 72 measurement days. Annually averaged R_s_ in 2024 (3.10 ± 1.86 CO_2_ µmol m^-2^ s^-1^) was higher than in 2023 (2.74 ± 1.89 CO_2_ µmol m^-2^ s^-1^), while the highest daily average R_s_ over the whole measurement period was reached on 23^th^ June 2023 (7.74 ± 2.0 µmol CO_2_ m^-2^ s^-1^). In 2024 the highest daily averaged R_s_ on 11^th^ July was lower by almost 1 µmol CO_2_ m^-2^ s^-1^ (6.83 ± 2.44 µmol CO_2_ m^-2^ s^-1^). We identified strong seasonal fluctuations with peak R_s_ during summertime with only small differences between the two years, but high seasonal differences (Figure 1). In 2023 annual precipitation was 37 mm below the long year average, while in 2024 it was 106 mm above. Only in summer the seasonal mean of VP_cur_ revealed an interannual variation, while the VPD showed variations throughout all seasons. Also, the determined Q_10_ values (Equation 5), using the daily average R_s_ and T_s_, revealed seasonal variability with the highest Q_10_ for both winter seasons in both years. However, the highest Q_10_ during winter 2023 exceeded the Q_10_ in winter 2024 by a factor of 2.6. In 2023 the Q_10_ continuously decreased till autumn, whereas in 2024, the lowest Q_10_ was reached in the summer season. The Q_10_ based on values throughout the whole year was similar with 3.32 ± 1.22 and 3.11 ± 1.12 in 2023 and 2024, respectively. Correlation tests separated by season (Table 2) revealed different responses of R_s_ in relation to the meteorological and microclimatic predictors. The aggregated values from 7 days prior to the actual measurement day (compared to 1-6 days) reflected the highest correlations consistently, which is why we focused only on these values. The lowest correlations and even negative correlations were present during summer for all parameters except for SWC and the summarized precipitation. In fact, during summer season 2023, the only significant positive correlation was found with the summarized precipitation 7 days prior to the measurement day.

**Figure 1:**
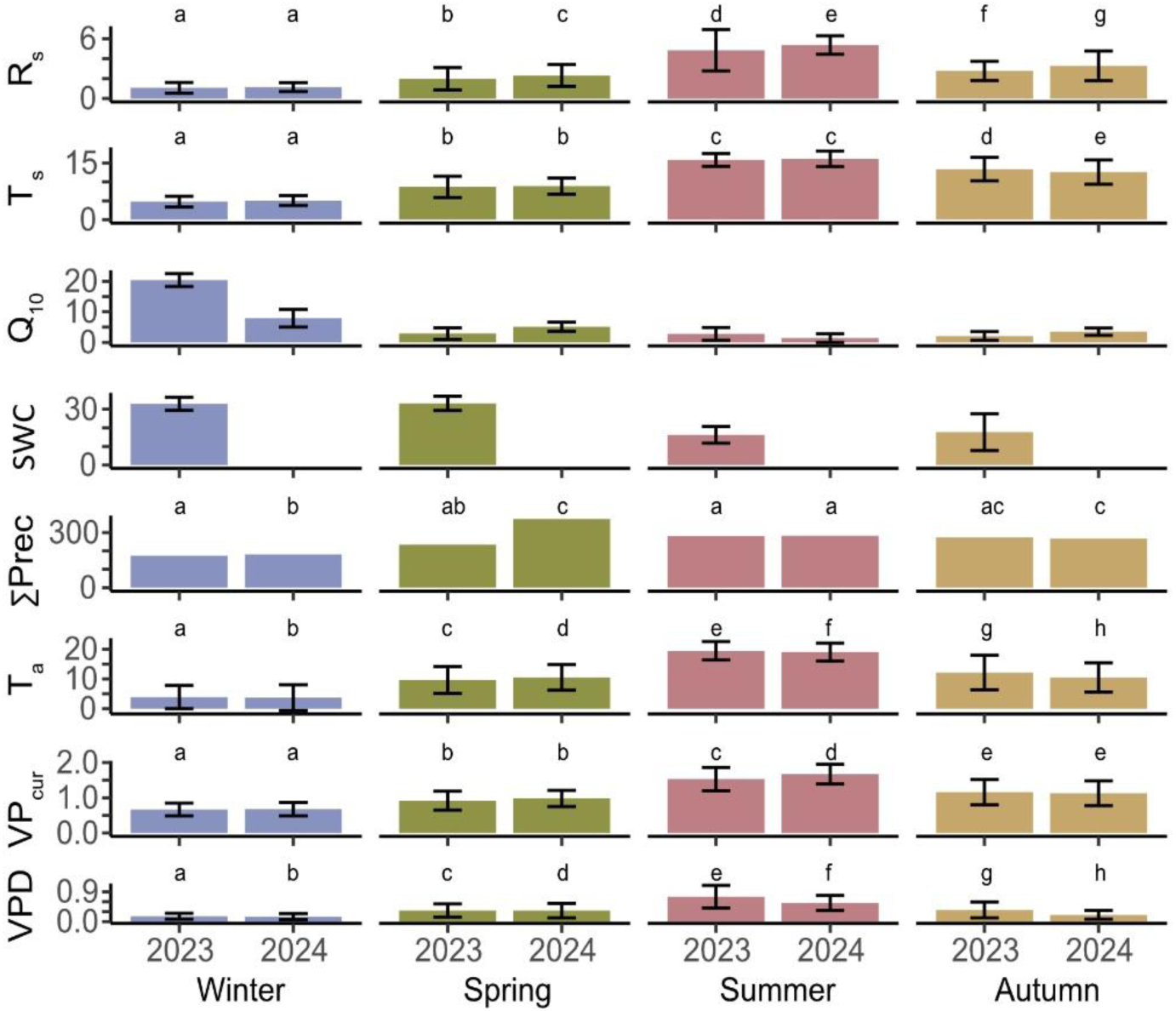
Seasonally averaged soil respiration (R_s_ [µmol CO_2_ m^-2^ s^-1^]), soil temperature (T_s_ [°C]), Q_10_, summarised precipitation (∑Prec [mm season^-1^]), air temperature (T_a_ [°C]), current vapour pressure (VP_cur_ [kPa]) and vapour pressure deficit (VPD [kPa]).

**Table 2:**
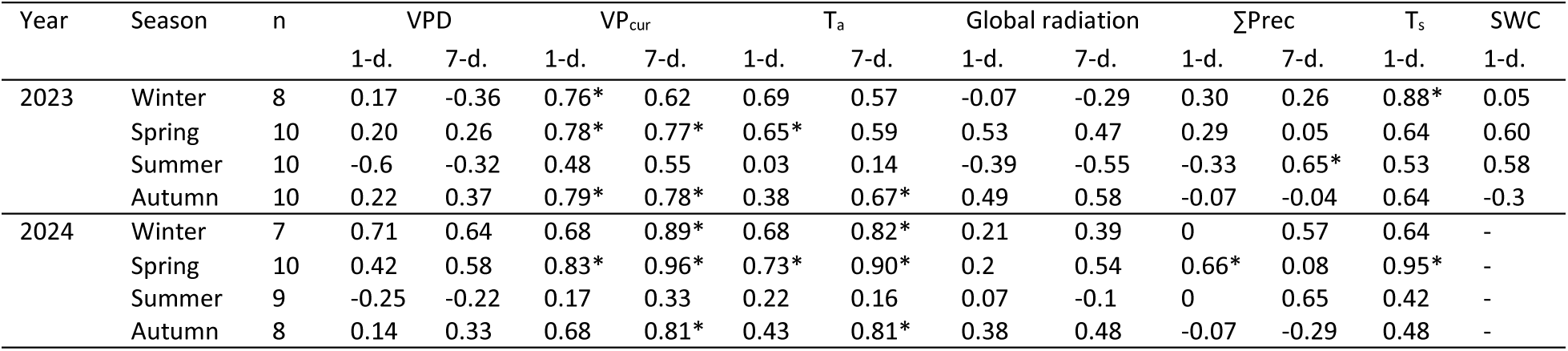
Spearman-correlation coefficients between daily averaged R_s_ and potential predictors as means value of the measurement day (1-d.) and the aggregated values seven day prior to the measurement day (7-d.). Meteorological predictors are vapour pressure deficit (VPD), current vapour pressure (VP_cur_), air temperature (T_a_), global radiation, sum of precipitation (∑Prec), soil temperature (T_s_) and soil water content (SWC). Significant correlation coefficients are marked with * (p-value < 0.05).

| Year | Season | n | VPD | | $VP_{cur}$ | | $T_a$ | | Global radiation | | $\Sigma Prec$ | | $T_s$ | SWC |
| --- | --- | --- | --- | --- | --- | --- | --- | --- | --- | --- | --- | --- | --- | --- |
|  |  |  | 1-d. | 7-d. | 1-d. | 7-d. | 1-d. | 7-d. | 1-d. | 7-d. | 1-d. | 7-d. | 1-d. | 1-d. |
| 2023 | Winter | 8 | 0.17 | -0.36 | 0.76* | 0.62 | 0.69 | 0.57 | -0.07 | -0.29 | 0.30 | 0.26 | 0.88* | 0.05 |
|  | Spring | 10 | 0.20 | 0.26 | 0.78* | 0.77* | 0.65* | 0.59 | 0.53 | 0.47 | 0.29 | 0.05 | 0.64 | 0.60 |
|  | Summer | 10 | -0.6 | -0.32 | 0.48 | 0.55 | 0.03 | 0.14 | -0.39 | -0.55 | -0.33 | 0.65* | 0.53 | 0.58 |
|  | Autumn | 10 | 0.22 | 0.37 | 0.79* | 0.78* | 0.38 | 0.67* | 0.49 | 0.58 | -0.07 | -0.04 | 0.64 | -0.3 |
| 2024 | Winter | 7 | 0.71 | 0.64 | 0.68 | 0.89* | 0.68 | 0.82* | 0.21 | 0.39 | 0 | 0.57 | 0.64 | - |
|  | Spring | 10 | 0.42 | 0.58 | 0.83* | 0.96* | 0.73* | 0.90* | 0.2 | 0.54 | 0.66* | 0.08 | 0.95* | - |
|  | Summer | 9 | -0.25 | -0.22 | 0.17 | 0.33 | 0.22 | 0.16 | 0.07 | -0.1 | 0 | 0.65 | 0.42 | - |
|  | Autumn | 8 | 0.14 | 0.33 | 0.68 | 0.81* | 0.43 | 0.81* | 0.38 | 0.48 | -0.07 | -0.29 | 0.48 | - |

### 3.2. Hot moments of R_s_

During the transition between spring and summer in 2023 (mid of May – mid of June) average R_s_ decreased to only 3 µmol CO_2_ m^-2^ s^-1^. This phase was characterised by increasing T_s_ and T_a_ but a lack of precipitation (Figure 2, A). Only a low rainfall event occurred, causing a slight increase in R_s_ by 0.72 µmol CO_2_ m^-2^ s^-1^, but a further decrease in the following week. During this whole phase, mean SWC dropped by 24 %_vol_. With an onset of rainfall with a total of 29.5 mm (approx. 3 % of 30-year average annual precipitation) a sharp increase in R_s_ by 6.3 µmol CO_2_ m^-2^ s^-1^ was measured, while average T_s_ and SWC increased only slightly by 1.5 °C and 1.3 %_vol_. The day after the strong rainfall event represented the highest daily R_s_ (7.74 ± 2.0 µmol CO_2_ m^-2^ s^-1^) over the whole measurement period. Additionally, VPD increased during the phase by 0.94 kPa but decreased directly afterwards the rainfall event by 0.33 kPa. During the phase the maximum average VPD was 1.37 kPa and R_s_ revealed negative correlation to T_s_ and VPD (ρ_s_ = -0.77 and ρ_s_ = -0.94, respectively), but positive correlation to SWC (ρ_s_ = 0.83). After the stronger rainfall event mean R_s_ dropped to 4.4 µmol CO_2_ m^-2^ s^-1^ just six days later to a level comparable prior to the strong decline. A comparison to the same time period in 2024 with more frequent precipitation revealed weaker fluctuations of R_s_ as in the previous year 2023 (Figure 2, B). However, we also identified additional phases characterized by low precipitation and low SWC i.e. during the end of August 2023 which was not as prolonged as the aforementioned phase. In comparison to VPD, we identified around 10 additional but shorter time periods in which VPD and R_s_ were correlated negatively. Interestingly, the two highest R_s_ rates over the whole measurement period were captured in 2023.

**Figure 2:**
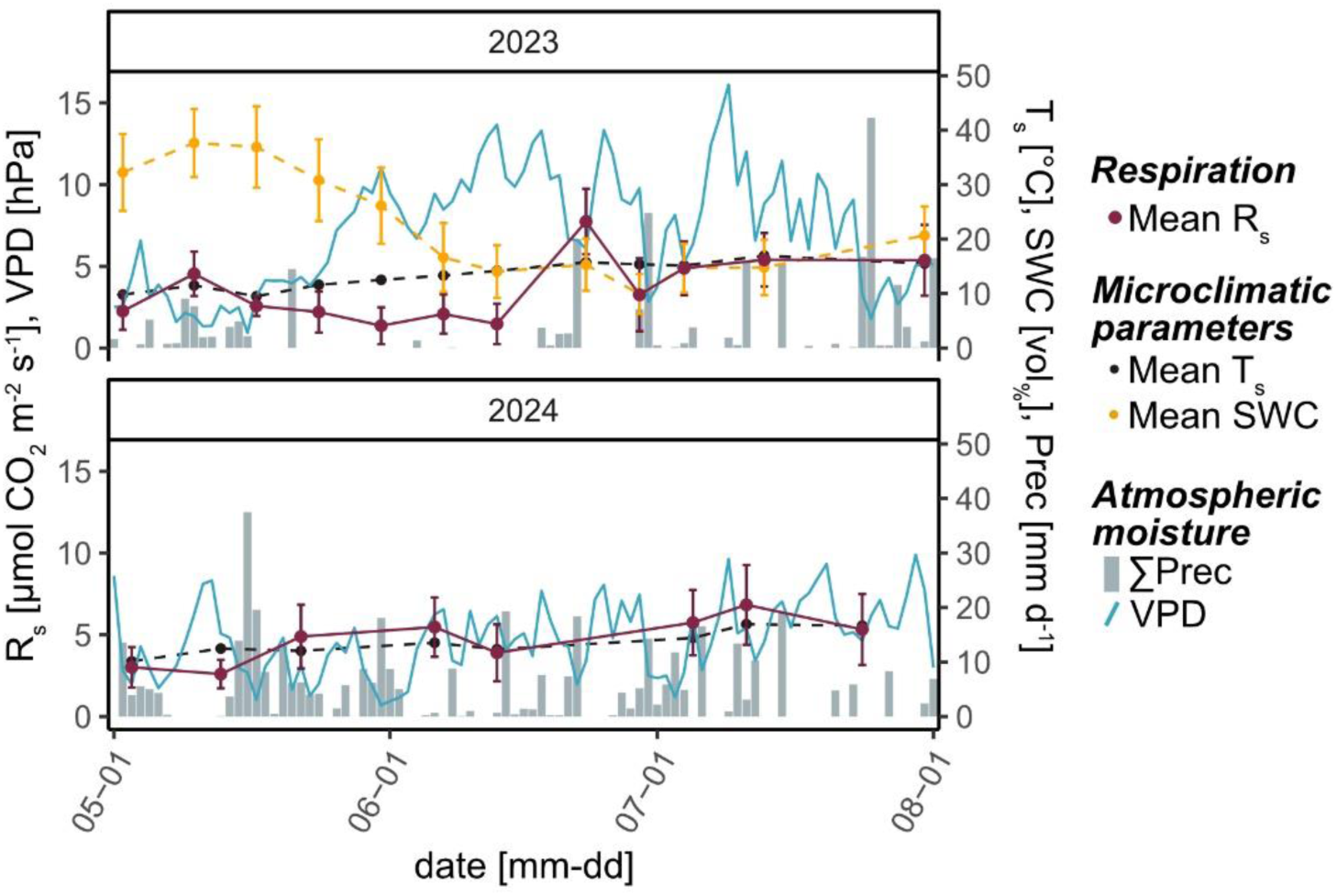
Excerpt of the timeseries from May to June for both observation years. Displayed are daily averages of soil respiration (R_s_), soil temperature (T_s_), soil water content (SWC), vapour pressure deficit (VPD) and summed precipitation (∑Prec). Note: VPD is displayed in hPa for better comparison.

### 3.3. Regression models with meteorological and microclimatic parameters

Individual linear regression models were used to determine predictors, that explain a considerable amount in temporal variability in R_s_ (Table 3). As predictor variables we used the downscaled meteorological data in comparison to locally measured values on plot level (T_s_, SWC). The highest explanatory contribution yielded the 7-day mean of the VP_cur_ from the HYRAS model, followed by the current VP_cur_, both clearly higher than T_S_, as the local variable with the highest explanatory yield. Multiple regression was used to combine predictors to the highest explanation in variability. Residuals of the models are reported as the quotient between modelled and observed R_s_. Values above 1 represented an overestimation and below 1 an underestimation (Figure 3). Absolute residuals are presented in Figure S5 (a-d) and overall model performance in Table S3. The four multiple regression approaches with the highest explanatory yield were (Formulas implemented in Figure 3 A-D):

**Table 3:**
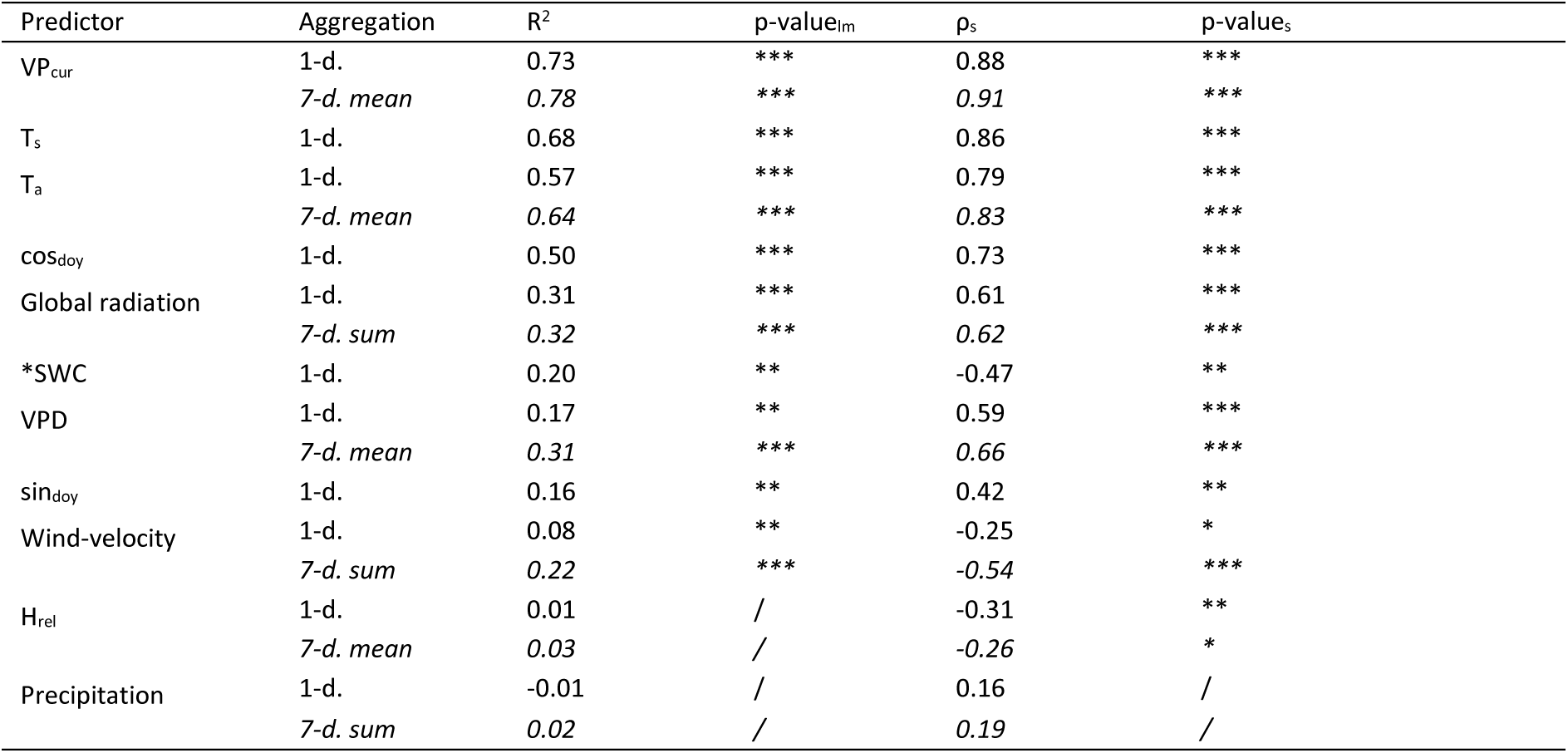
Results of individual linear models with and without temporal aggregation of meteorological parameters. Explanation of variability (R^2^) in R_s_ determined with individual linear models and correlation coefficients (ρ_s_) for meteorological predictors with significance of linear regression (p-value_lm_) and correlation tests (p-value_s_). Significance levels are: / ≥ 0.05, * < 0.05, ** < 0.01, *** < 0.0001. Correlation coefficients between the predictors itself are listed in the supplemental materials (Figure S1).

A. *Exponential T_s_ and SWC model (R^2^= 72%)*: The exponential fit of T_s_ and SWC (Equation 4) resulted in the second highest explanation in variability. The maximum deviation occurred during the transition between spring and summer (compare chapter 3.2.) which reflects the highest deviation for this model.
B. *Linear T_s_ model (R^2^ = 68%)*: The linear fit (Table 3) showed three strong peaks of overestimating the actual R_s_ by a factor of > 2. Similar to the previously applied model (A.) the highest deviation was present during the drought period (compare chapter 3.2.) and the following rewetting which resulted in a strong decline and a residual factor close to 0.5 which reflects the strong underestimation of this model. However, the third peak in deviation during autumn is not visible in the previously applied model (A.). In order to better compare the results with model A, the fit still revealed a lower explanation in variability (60%) if modelled only for 2023.
C. *Exponential T_s_ model (R^2^ = 68%)* The exponential model fit (Equation 3) described a similar proportion in variability as the linear fit (Model B). However, peaks occurred more often and actual R_s_ was overestimated five times by a factor > 2. This model showed a contrast in estimations to the linear fit with unspecified over- and underestimation while the linear fitted model underestimated R_s_ systematically during winter 2023 – 2024. The deviation of this model during the end of 2024 is also vice versa as in the prescribed model (B.). In order to better compare the results with model A, the fit revealed the lowest explanation in variability (58%) if modelled only for 2023.
D. *Meteorological model* (*R^2^ = 0.84*): The highest explanation in variability of R_s_ is represented by the 7-day-mean VP_cur_ (Table 3). T_s_ was not included into this model due to high co-linearity with the VP_cur_ and the 7-day mean VP_cur_. The VP_cur_ also includes T_a_ as a variable and was therefore not included as a separate predictor. As the next highest predictor, the cos_doy_ was included. The residuals of this model contained high information content and were systematically explainable in all seasons by positive relationship with VPD (Figure S3. Since VPD showed a high collinearity with the VP_cur_, we used the detrended residuals of the VPD as a parameter in Model D (Figure S4). These residuals represented low VPD (rather wet atmospheric conditions) as negative value, and high VPD (rather dry atmospheric conditions) as positive value. Including the new predictor increased explanation in variability by 6 %. The residual factor of this model exceeded, similar to the model (A.), the actual R_s_ value by a residual factor of > 2 only once, but reflects in general the lowest residual factors. In order to better compare the results with model A, the fit still revealed the highest explanation in variability (77%) if modelled only for 2023.

**Figure 3:**
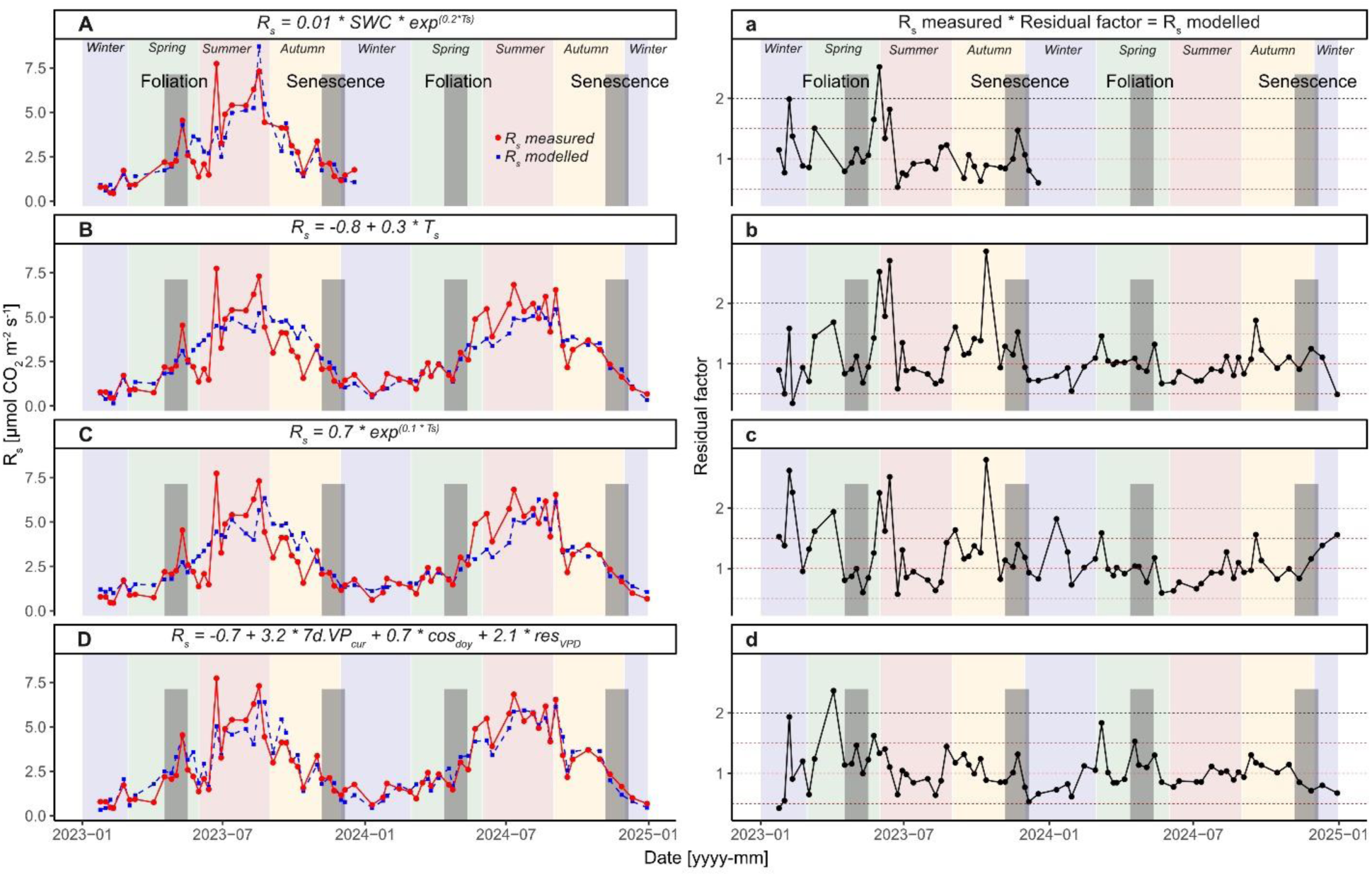
Modelled soil respiration (R_s_) in comparison to measured R_s_ (A-D) and residual factor of the models (a-d) for the whole measurement period. Colored areas represent seasons. Note: R_s_ is only modelled for the respective measurement day.

Again, in the multiple regression, the meteorological data and the season showed the best explanatory power. All models presented an underestimation of R_s_ during summer 2023 but an overestimation during autumn and the senescence in 2023. Surprisingly, especially during the “hot moment” (compare chapter 3.2.), the *meteorological model* (Figure 3 D) using the downscaled meteorological data performed best and presents the lowest deviations from the observed R_s_. However also the model using the locally measured data (T_s_, SWC) (Figure 3 A) showed a good prediction quality and followed the measurements with only little deviation. During spring 2023 the *meteorological model* did not perform better and presents the largest deviation compared to the other three models. Commonly for all models, a lower deviation and better performance is reflected during 2024 which was not characterized by a drought period.

### 3.4. Spatial predictors – hot spots of R_s_ and the influence of forest structure

Spatial variability in R_s_ within the ECOSENSE Forest was high and hot and cold spots occurred during all seasons. As a hot spot we defined plots with significantly higher respiration compared to the average over the measurement area while cold spots were defined as plots with significantly lower R_s_ (Figure 4). Especially one plot was always considered a hot spot (marked with a red point), whereas all other hot and cold spots occurred only in specific seasons. The patterns stayed almost persistent throughout both measurement years. However, an analysis of hot and cold spots based on structural parameters did not reveal any clear and significant patterns. The overall occurrence of hot and cold spots is rather low, especially with regard to the different tree species. The persistent hot spot was characterized by a large number of deciduous trees (10) occurring within the 5 m buffer zone. The two hot spots in spring were associated with a high number of trees within the 5 m buffer zone, whereas, conversely, a high number of trees formed cold spots in winter. The northern part with higher R_s_ is densely occupied by deciduous composition while the transect on the north-south axis mostly consists of coniferous patches, implying an important influence of tree species on R_s_. The species effect on R_s_ (Figure 5 A) was only significant during summer season with higher R_s_ associated with deciduous trees compared to coniferous and mixed composition.

**Figure 4:**
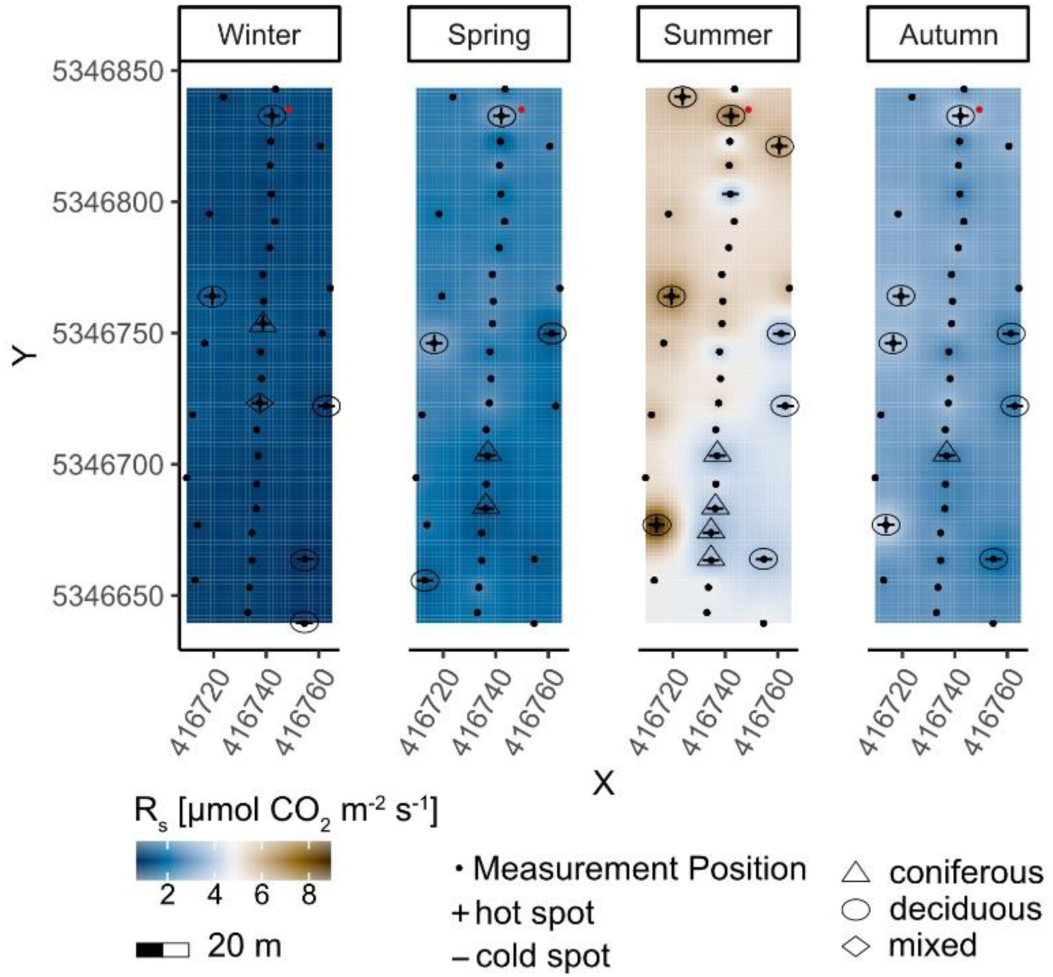
Spatial variability of soil respiration (R_s_) represented as an inverse-distance-weighted interpolation, separated by season for both observation years.

**Figure 5:**
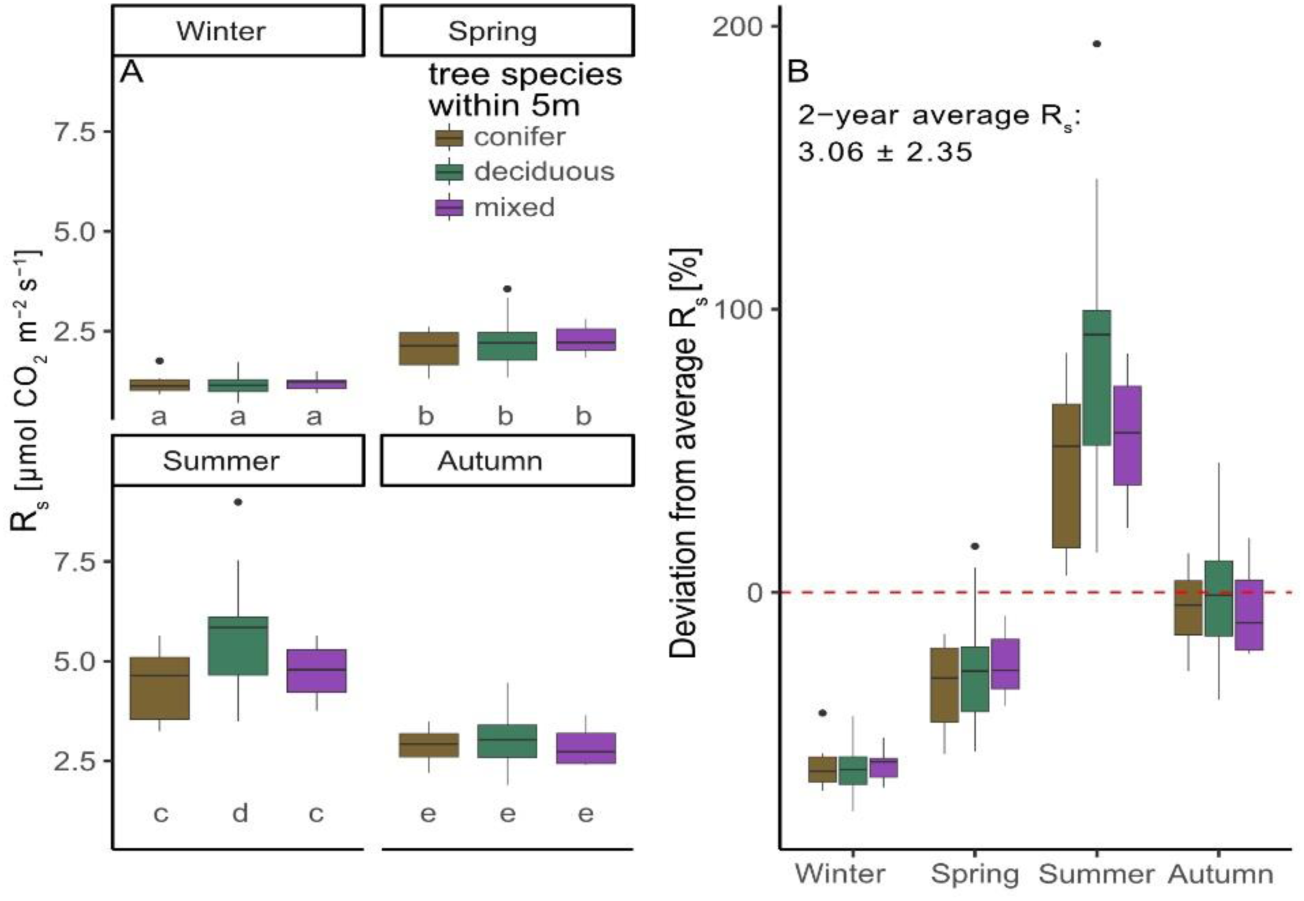
Soil respiration (R_s_) in dependence of tree species composition (A) and seasonal, plot-wise deviations from the mean R_s_ across both observation years (B). Significant differences between the groups are indicated by letters.

The deviation from the overall average R_s_ was least pronounced during the autumn season, with the slightest average deviation for deciduous composition while winter and spring deviations were below average R_s_ (Figure 5 B). Interestingly, in average, measurements during summer season exceeded the annual average R_s_ up to almost 100% in deciduous composition, while deviated by almost 200%.

Unlike tree species composition, the other forest stock features did not show clear influences (Figure S6) on R_s_ with exception of a significant negative correlation between tree height of conifers in spring, summer, and autumn and a similar effect of DBH. Within the mixed composition, correlations with tree height tended to be positive. Notably strong negative correlations were also observed with the distance to tree(s) during the vegetative seasons within almost all compositions. Overall, within the deciduous composition, correlations were the weakest but especially regarding the DBH, correlations changed from negative during winter and spring to positive during summer and autumn. In general, the influence of the season on the results becomes evident. The categorized distance (compare chapter 2.4) to the nearest tree trunk (Figure 6) revealed constantly higher R_s_ closer to the trunk while this effect was in tendency vice versa for T_s_ and SWC.

**Figure 6:**
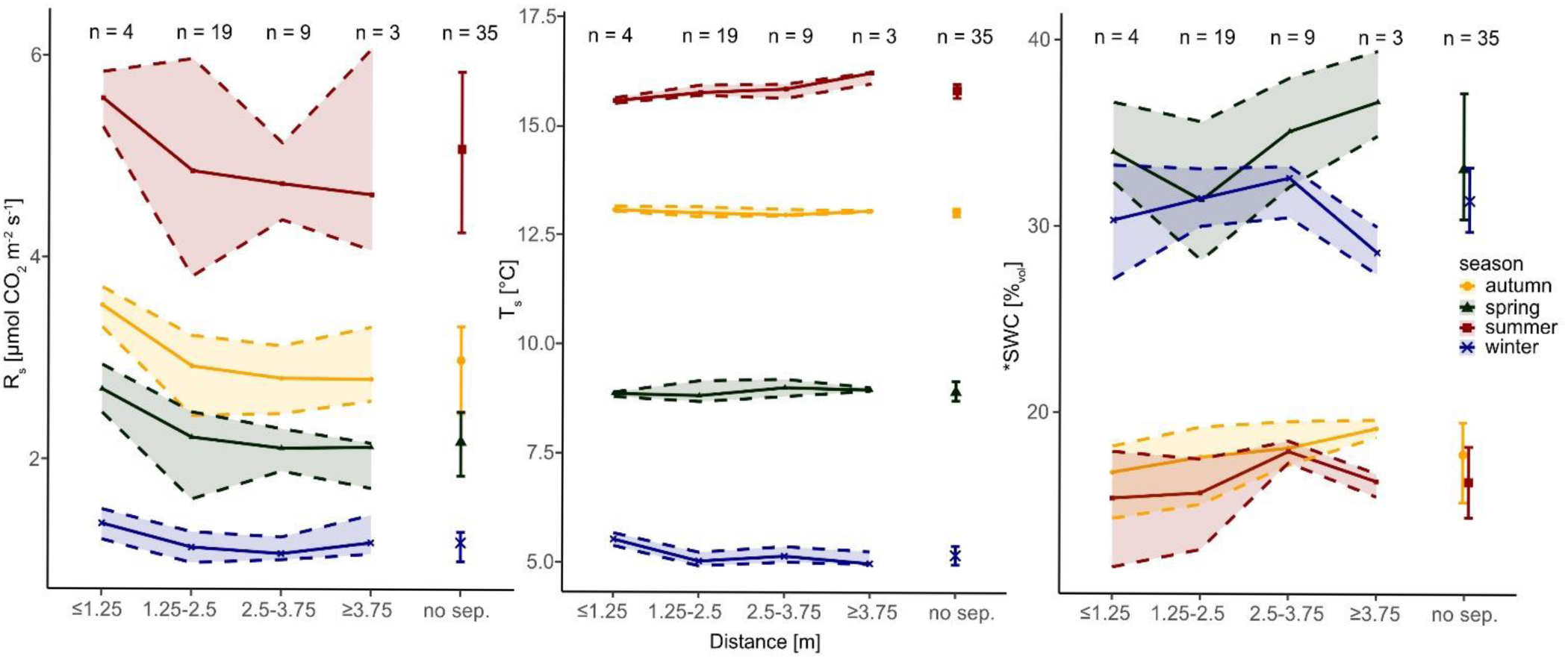
Soil respiration (R_s_), soil temperature (T_s_) and soil water content (SWC) in dependence on distance to the nearest neighbouring tree and season. Values represent averages per plot for the corresponding value. Shaded area and error bars represent the 75% and 25% Quantiles. Values are based on the total amount of measurements (n=2295). *SWC was measured only in 2023.

## 4. Discussion

### 4.1. Temporal patterns in R_s_ are controlled by meteorological conditions prior to measurements

The basic hypothesis of our study was the central role of meteorological parameters as controls of soil respiration (R_s_). This could be confirmed in several aspects. Two meteorological parameters and additionally the summer-winter contrast showed a superior explanatory power compared to locally measured soil temperature (T_s_) and soil moisture (SWC). It could also be confirmed that up to 7-day time lagged meteorological data, which were taken as a downscaled feature from a large scaled meteorological model, can better explain R_s_ than only the current observations alone lets us suggest and confirm, that R_s_ of the observed forest reacts rather sluggish to environmental changes (Ekblad et al., 2005; Kuzyakov and Gavrichkova, 2010). VP_cur_ combines exponential air temperature (T_a_) and relative humidity (H_rel_) in one variable. VPD as well acts on the whole ecosystem including the canopies of the trees, and by this way also directly on the physiological state of the trees including root respiration (compare also chapter 4.2). Its role as a controlling variable of R_s_ has not been described so far. The positive relationship indicated the importance not only of temperature, but also atmospheric water for temporal R_s_ patterns. In the following, we discuss possible effects of atmospheric moisture conditions on R_s_.

Only one study addressed explicitly the effect of air humidity (H_rel_) on R_s_ (Kukumägi et al., 2014). In a hemiboreal forest ecosystem, consisting of young, planted silver birch, the authors demonstrated in the second year after the application of air moisturising (7% increase in H_rel_, which increases the VP_cur_ if the T_a_ stays the same) decreased R_s_ rates which can be seen as contrast to our results. However, the authors demonstrated additionally an increase in SWC compared to ambient conditions. While a certain level of SWC reflects beneficial processes on microbial processes as well as tree water supply, excessively high SWC can promote anaerobic conditions and thus limit the supply of oxygen in the rhizosphere and have an inhibitive effect (Skopp et al., 1990). However, in the same experimental setup, Kupper et al. (2011) determined lower sap-flux and transpiration of the saplings even on days without moisturising. These are changes which also affect carbon fluxes and might influence R_s_ in longer term. However, as Kukumägi et al. (2014) demonstrated, that the decrease in R_s_ could potentially be explained rather by investments in root growth of the understorey vegetation. In the ECOSENSE Forest, understorey is only sparsely present and trees are much higher and of older age, and therefore results not overall comparable. The results of our study compared to the artificially created increase in H_rel_ suggest that there might be an optimal level of H_rel_ (i.e. VP_cur_) that promotes R_s_ and is beneficial for ecosystem activity.

In the present study we also point out to a strong correlation and co-linearity between the VP_cur_ and T_s_, which might be a potential reason why explanations in variability of both models were strong. Beside the effect that the VP_cur_ was not tested in most other R_s_ models, T_s_ might be in other research layouts a more suitable predictor for R_s_ for possibly two reasons: firstly, we covered a relatively large area of around one hectare with a high density in measurement positions. This usually results in a high level of background noise of local measured soil parameters, with greater variability than on smaller areas. Secondly, reference depths are variable between different studies. We measured T_s_ in 13 cm depth, while e.g. Moyano et al. (2008) found highest correlation with T_s_ in 2 cm depth. Ruehr et al. (2010) suggested T_s_ measurements within 1-5 cm depth for lag estimations, but presented for depths of 10 cm an exceptionally high Q_10_ which cannot be supported by our findings of Q_10_ values. The values determined in our study were similar to the Q_10_ determined by Khomik et al. (2006) from 2 cm soil depth. We also calculated seasonally separated Q_10_ values as suggested by Khomik et al. (2006) with the limitation that a T_s_ range of up to 10 °C within one season might be unrealistic. The higher Q_10_ values during wintertime and lower Q_10_ during summertime suggest a shift from a temperature-controlled to a climate/moisture-controlled regime in summer. Additionally, the change in correlations between summer and winter season underlined our assumption, as correlation between R_s_ and SWC and the summarized precipitation was higher during the spring and summer season, but reversed for the other predictors. A similar change in controls in R_s_ was also suggested by Hereș et al. (2021). As T_s_ was only measured in combination with R_s_, we cannot make any more precise statements about lag-effects of T_s_ on R_s,_ as e.g. Ruehr et al. (2010). However, we also point out to a strong correlation between T_s_ and global radiation and the summarised global radiation and therefore highlight the effect of global radiation on T_s_ development (Kuzyakov and Gavrichkova, 2010).

Even though the *meteorological model* performed well, the absolute residuals (Figure S5) were considerably high during the vegetation season compared to non-vegetative period. During wintertime, the ecosystem stays dormant due to lower temperatures, short daylight and the absence of foliage of deciduous trees and absolute R_s_ is low. Photosynthetic activity is at rest except for coniferous evergreen trees (Hansen et al., 1997; Jochheim et al., 2022) and due to lower T_s_ and low nutrient availability, microbial activity as well. Photosynthetic activity with its peak during the summer drives R_a_ and therefore the total R_s_ (Kuzyakov and Gavrichkova, 2010; Tang et al., 2005a). Microbial activity is also higher during the summer season, because enzymatic activity depends on T_S_ (Schindlbacher et al., 2011; Schindlbacher et al., 2012) but also on root exudate availability, which in turn is driven by photosynthesis (Kuzyakov and Gavrichkova, 2010). Reduced correlations with most of the presented predictors including local parameters during the vegetation period further underline the importance of the phenology on temporal R_s_ patterns. Yet for this research solar radiation is the only available predictor which might be used as a proxy for photosynthetic activity, beside the manual observation of leaf developmental stage. The latter resulted in a direct increase in R_s_ due to foliation and decrease in R_s_ due to senescence (Figure 3). Therefore, we implemented smooth seasonal shifts with the cosine angle function cos_doy_ in our model, which created a peak during summertime. This obviously still lacks the true representation of the phenology. The overestimation, during spring visible in all presented models, could be due to the fact that trees invest sugars into biomass at first (Kuzyakov and Gavrichkova, 2010) and therefore R_s_ does not follow the steady increase of the predictors. Furthermore, R_h_ might be low due to low exudation of easily available substrates in the rhizosphere. However, the *meteorological model* created a broader range in residuals during summer 2023 compared to summer 2024 which can be attributed to the rather dry conditions. During this dry period the model is not capable in representing the high variability in responses to rewetting after rainfall. This suggests a strong spatial and individual response on plot-level. The widely applied models using T_s_ as the only predictor or in combination with SWC, especially during extreme events in R_s_, did not deliver the same performance as the presented *meteorological model*. The simple use of T_s_ as predictor demonstrates that, in periods where SWC is the limiting factor, these models do not provide a sufficient performance (Ruehr et al., 2010) and alternative predictors are of great importance, especially in the scope of an ongoing climate change. We suggest that the physiological activity influencing the R_s_, is better reflected by the smoothed atmospheric conditions than by soil parameters. Models for R_s_ still lack in the true representation of influences of phenology on R_s_ and the influences of carbohydrate supply (Mitra et al., 2019), therefore the VP_cur_ might represent a valuable predictor for plant physiological activity.

### 4.2. Collapse of R_s_ and increase due to rewetting

In our second hypothesis we stated that R_s_ pulses after rewetting might be a hot moment contributing relevantly to total R_s_. We were able to clearly identify one rewetting period as an actual “hot moment” of R_s_ which was the highest measured daily R_s_ over the two-year investigation period. During the drying period, the usual patterns of positive correlation between T_s_ and R_s_ were reversed due to a decline in SWC. The following pattern of a subsequent rewetting pulse of R_s_ has already been reported in the literature (Birch, 1958; Jochheim et al., 2022; Ruehr et al., 2010; Tang et al., 2005b). However, there are also contrary results. Schindlbacher et al. (2012) observed a decline due to drought and a regeneration of R_s_ due to rewetting, but not a peak. The authors attributed the lack of this effect to the importance in homogeneous rewetting of the forest floor, which might not have occurred due to preferential flow paths created by drought induced hydrophobicity (Schindlbacher et al., 2012). Overall, the data obtained confirm our hypothesis and we suggest similar underlying mechanisms to those already reported (e.g. Birch, 1958), that the strong increase was an effect of reactivation of microbial activity. Jochheim et al. (2022) additionally suggested a potential collapse in R_a_ associated with *Fagus sylvatica* due to a stronger decline associated with these species, which cannot be supported by our data, as all plots were affected similarly. However, we cannot exclude that R_a_ was not affected by the drought period. The vapour pressure deficit VPD can also be used as an estimate for aboveground plant stress, as trees open or close their stomata when VPD is low or high, respectively (McAdam and Brodribb, 2015). In comparison, the summer and autumn of 2023 showed a higher atmospheric VPD than 2024, indicating drier aboveground conditions. Sanginés de Cárcer et al. (2018) showed the effects of species specific tree growth reduction for VPD values above 1.5 kPa, decoupled from soil water limitations. This value can be used as an estimate at which the stomata of *Fagus sylvatica* (Lendzion and Leuschner, 2008) and *Pseudotsuga menziesii* trees (Meinzer, 1982) reduce conductance, which affects carbon uptake and therefore also carbon allocation and respiration by tree roots (Hsiao, 1973). In our study, there was only one day within the two years, which represented a VPD > 1.5 kPa. This might be due to the type of calculation of VPD using the daily averaged values. The VPD is therefore smoothed and represents only the average for the corresponding day, neglecting peaks. If the VPD is calculated using the maximum T_a_, values above 1.5 kPa were 4 times more common in 2023 compared to 2024 and the aforementioned hot moment is particularly affected by the high VPD. Interestingly, we detected a higher average R_s_ in 2024 compared to 2023, but the two highest R_s_ rates were reached in 2023, correlating to the precipitation deficit within the ecosystem and reflected ecosystem stress within the soil. This might also be reasoned in the different precipitation patterns during both years, with larger gaps in 2023 (Figure S2). Our results highlight the importance of temporally high-resolution measurements to capture these specific hot moments, since an interpolation of R_s_ during these phases might lead to false conclusions. The strongly differing correlations of VPD and R_s_ during the individual seasons underline the VPD as an important factor for R_s_.

### 4.3. Forest structural features and hot spots in the ECOSENSE Forest

In agreement with our hypothesis, we observed larger R_s_ for deciduous than for coniferous patches during summer. However, tree species did not affect R_s_ in winter, which is in contrast to Jochheim et al. (2022) and our hypothesis. A generally higher R_s_ for broadleaf species compared to coniferous species was also reported in the literature (Raich and Tufekciogul, 2000). Longdoz et al. (2000) presented differences in R_s_ associated especially with *Fagus sylvatica* and *Pseudotsuga menziesii* patches in a slightly different climate. They underline differences in fresh leaf litter and lower C:N and lignin:N as indicators for decomposition as possible reasons for these patterns. Another reason might be the difference in transport processes through phloem, which was summarized by Kuzyakov and Gavrichkova (2010), to be slower for coniferous species compared to the group of angiosperms, which includes *Fagus sylvatica*. Furthermore, *Pseudotsuga menziesii* forests reflected root biomass increase in earlier stages but stagnation to decrease at later stages of competitive growth after forest canopy closure (Vogt et al., 1987). The ECOSENSE Forest shows a dense canopy closure during summer with only sparse midstory. This might indicate rather investments in overstorey biomass growth.

The differences in tree species also seem to reflect the spatial pattern in the ECOSENSE Forest, as abundance of *Fagus sylvatica* is higher in the northern part of the area. Especially during the summer season, we detected a larger variability compared to the winter season which could also be reasoned in the photosynthetic activity of trees within the forest. Regarding the strong heterogeneous and seasonal pattern, surprisingly, the deviation in R_s_ during autumn season from the average over the two-year measurement period was lowest and not as variable between the species composition as during summer. Fresh leaf litter combined with water availability and average T_s_ during fall might promote microbial activity and thus raise R_h_ to a higher level compared to spring in temperate forest ecosystems. Simultaneously, R_a_ is reduced due to senescence. This also reflects the smooth transition from dormancy to activity and vice versa. The results suggest that R_s_ during autumn could reflect the average annual R_s_ in a temperate forest best, even though the forest reflects very heterogeneous stock features. Nevertheless, this highlights the importance of studies conducted over lager spatio-temporal scales. Short measurement periods or campaigns can not capture the whole seasonality and might bias the estimates of annual R_s_. Conclusions regarding upscaling should therefore be reviewed critically, not only on spatial (Hereș et al., 2021) but also on temporal scale.

We can partially confirm the hypothesis that R_s_ is higher close to the trunk. We highlighted a negative effect of the distance to the nearest tree, although the categorization of distances was more conclusive than the direct correlations, even though the latter were rather negative. Especially the varying strength during the seasons suggest a corresponding contribution of R_a_ to the measured R_s_ since neither T_s_ nor SWC clearly explained these patterns. The distance to the tree trunks has been rarely studied, but available studies explain the observed higher root biomass closer to the trunk (Saiz et al., 2006b) or differences in forest floor thickness (Scott-Denton, 2003). Also Reynolds (1970) reported a negative relation between root length and the distance to trees for *Pseudotsuga taxifolia*. Jochheim et al. (2022) report similar patterns in R_s_ in comparable distances as our categorised approach. The results indicate the strong influence of R_a_ on total R_s_ due to the higher root biomass closer to trunks especially due to the strong decline from < 1.25 m to 1.25 - 2.5 m distance. This steep decline closer to the stem compared to positions further away from the stem were also shown by Saiz et al. (2006b) but with much finer resolution. The results together can also indicate physical effects. CO_2_ produced below the tree, coupled with higher root biomass close to the stem, cannot diffuse through the trunk and therefore might diffuse from the soil close to the base of the tree stem. Due to our approach of a structured measurement grid and the precision range of ± 1 m, we are not fully able to prove this hypothesis. Intensive measurements close to stems need to be coupled with soil biological analysis (root density, microbial biomass) to prove the underlying processes for stemfoot R_s_ gradients.

Surprisingly, the tree height revealed negative relationship with R_s_, which was significant for coniferous composition especially during the vegetative seasons. To our knowledge, effects of tree height on R_s_ have not been reported in the literature until now. This effect could partly be explained by a longer pathways and in general slower transport of assimilates within coniferous trees to reach the rhizosphere (Kuzyakov and Gavrichkova, 2010), which could result in even more time-lagged R_s_. This is in contrast to results reported by Mitra et al. (2019), who did not find any relationship between the height of trees and the lag in R_s_. In order to contextualize the effect of tree height on R_s_ within existing research, stand age or DBH could be used. Trees in our research area tended to show larger DBH with larger tree height. Søe and Buchmann (2005) presented positive relationship between R_s_ and the DBH within 4 m distance, a result that only partially aligns with our findings. Saiz et al. (2006a) suggest that R_a_ contributions are higher in younger stands compared to older stands connected to higher root activity and productivity in younger Sitka spruce (*Picea sitchensis*) stands. Similarly, Klopatek (2002) reported lower R_s_ associated with older aged Douglas fir stands, related to lower fine root productivity even though fine root biomass was higher compared to the younger aged stand. These results might be related to the previously mentioned competition among the coniferous species, tending to prioritize vertical growth. The complex relationships between tree age and associated tree height, DBH, root biomass and root length as well as transport processes within trees are yet not fully explainable with our study design but the results indicate that tree height should be considered in future research for R_s_.

We also point out to a slightly higher soil-gas diffusivity in the northern part (Dystric Skeletic Cambisol). Differences between the soil types in regard to the other specified physical and chemical properties are not very pronounced except for a slightly higher soil-organic carbon content in the topsoil horizons, cation exchange capacity (CEC) and a broader C:N. However, soil properties could be very heterogeneous throughout forest soils (Stutz and Lang, 2023) and a clear distinction of soil types within the ECOSENSE Forest is yet to be determined. A transition zone could be located approximately in the center of the study area. Due to rather similar chemical and physical properties of the described soil profiles, we do not expect larger influences of soil properties on R_s_. Biological analyses such as root distribution and microbial biomass, chemical and physical properties of each of the 35 plots within the ECOSENSE Forest may support our results that tree species and their associated micro-habitat as root systems, substrate availability and microorganisms shape R_s._

## 5. Conclusion

Reliable large-scale soil respiration models are required to consider land use and its changes in climate research but the mostly site-specific models based on existing studies make an upscaling difficult. Our results show that the temporal pattern of soil respiration could clearly be explained better using downscaled meteorological data compared to the common local parameters soil temperature and soil moisture. Especially during extraordinary periods as the collapse in R_s_ during drought and the boost after rewetting were reflected with a lower residual error. These results from a mixed forest are unique and further research is required for generalization. However, interpolated meteorological data also offer the advantage of long, daily timeseries, which also offers possibilities to estimate R_s_ into past periods. We suggest, as upscaling approach, to remodel existing R_s_ timeseries with meteorological parameters and to include these parameters in future R_s_ studies. Additionally, our results provide important new ecological insights particularly regarding tree species composition and forest stock features. Our local R_s_ maps revealed places with very high or very low R_s_ independently from tree distribution. Yet, this could not fully be explained by the available parameters but were spatially consistent over two consecutive observation years. However, the small-scale local heterogeneity reflected the importance of non-meteorological parameters such as soil and vegetation. The contrast between deciduous and coniferous tree species had a significant contribution to explain the local variation of our data. Comprehensive meteorological data can help to describe the temporal pattens of R_s_, which is influenced by land-use specific spatial factors. These can be extracted from existing data and supplemented with specifically targeted studies to fill gaps.

## Acknowledgements

We thank the laboratory team of the Chair of Soil Ecology for supporting the laboratory analysis. Furthermore, we want to thank Luna Reimer, Chris Huck and Fiona Bauer for their support during the start of the measurement campaign. We also thank the team members of the Chair of Sensor-based Geoinformatics and the Chair of Forest Growth and Dendroecology who worked on the forest tree inventory and helped in field, digitizing and validating the dataset. We thank Kristin Steger for the internal pre-submission review.

## Data availability

The data from this research are deposited online on FreiData (https://doi.org/10.60493/hk8sq-c3c66). TLS scans can be accessed at 3Dtrees.earth (2025) under dataset ID 195. The meteorological data of HYRAS (DWD, 2025) and ERA5 (Copernicus Climate Change Service, 2019) can be accessed online.

## Statements and Declarations

### Author Contributions

Julian Brzozon: Writing – original draft, Investigation, Methodology, Formal analysis. Patricia Schwarzkopf: Investigation. Friederike Lang: Writing – review & editing. Teja Kattenborn: Writing – review & editing, Investigation, Methodology, Formal analysis. Julian Frey: Writing – review & editing, Investigation, Methodology, Formal analysis. Helmer Schack-Kirchner: Writing – review & editing, Methodology, Conceptualization, Funding acquisition.

## Abbreviations

CO2: carbon dioxide
Rs: soil respiration
Ts: soil temperature
SWC: soil water content
VPcur: current atmospheric vapour pressure
VPD: atmospheric vapour pressure deficit
Hrel: relative humidity
Ta: air temperature
DBH: diameter at breast height

