## Supplemental Material for "Predictions of spatio-temporal patterns of soil respiration in a mixed forest: Downscaled meteorological data outperform local parameters"

[1]Regional board Freiburg, Bissierstrasse 7, 79114 Freiburg, Germany

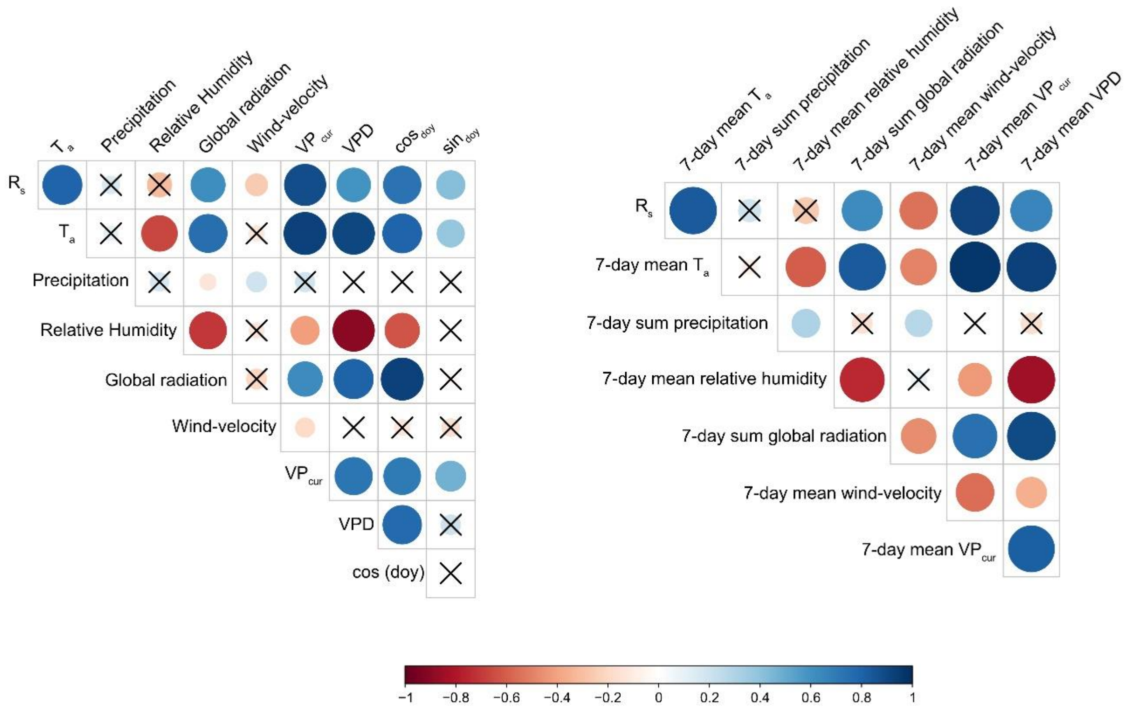


*Figure S1: Correlation matric between meteorological predictors and soil respiration ($R_s$) over the two- year measurement period.*

*Table S1: Selected research from European Forest Ecosystems covering soil respiration measurements and seasonality at least for 1 year.*

| Location | Study Area [~ha] | Number of measurement positions [n] | Measurements [days] | Campaign length [days] | Time coverage [years] | Reference |
|---|---|---|---|---|---|---|
| Germany | 0.5 | 36 & 144 moving | 27 | 1-3 | 2 | Søe and Buchmann (2005) |
| Switzerland | 4 | 17 & 1 automated | 36 continuous | 1 | 2 | Ruehr et al. (2010) |
| Switzerland | 4 | 10 | 36 | 1 | 2 | Ruehr and Buchmann (2010) |
| Czech Republic | 4 | 81 | 8 | 2 | 1 | Hereș et al. (2021) |
| Ireland | 0.4 | 120 | 11 | n.d. | 1 | Saiz et al. (2006) |
| Germany | 0.5 | 36 | 35 | 1 | 3 | Knohl et al. (2008) |
| France | 0.6 | 6 | 22 | 1 | 1.5 | Epron et al. (1999) |
| Germany | 1 | 35 | 72 | 1 | 2 | This study |

*Table S2: Methods and measurement devices for determination of soil chemical and physical properties.*

| Parameter | Method | Measurement device |
|---|---|---|
| pH | Gutachterausschuss Forstliche Analytik (2022) | Automatic sampler (905 Titrando, Metrohm, Germany) |
| C:N | Gutachterausschuss Forstliche Analytik (2022) | CHNS elemental analyzer (vario EL cube, Elementar Analysensysteme GmbH, Germany) |
| CEC | Trüby and Aldinger (1989) | ICP-OES (5800, Agilent, USA) |
| $D_s/D_0$ | Kühne et al. (2012), (Schack-Kirchner et al. 2001) | Micro gas chromatograph (CP2002, Chrompack, The Netherlands) |
| Particle Size | Hartge and Horn (1989) | |
| $\phi_{soil}$ | Hartge and Horn (1989) | Vacuum-Pycnometer |
| $\rho_{bulk}$ | Hartge and Horn (1989) | Vacuum-Pycnometer |

*Table S3: Model performance and model predictors (meteorological model). P-value levels are: / ≥ 0.05, * < 0.05, ** < 0.01, *** < 0.0001. Note: Model A was not heteroscedastic.*

| Model | Predictor | Estimate | Std. Error | p-value | Robust std. Error (HC3) | p-value | Breusch Godfrey p-value |
|---|---|---|---|---|---|---|---|
| A | SWC | 0.01 | 0.002 | *** | | | 0.93 / |
| | $T_s$ | 0.2 | 0.015 | *** | | | |
| B | Intercept | -0.75 | 0.32 | * | 0.20 | *** | 0.30 / |
| | $T_s$ | 0.33 | 0.03 | *** | 0.03 | *** | |
| C | a | 0.73 | 0.12 | *** | 0.05 | *** | 0.36 / |
| | $T_s$ | 0.12 | 0.01 | *** | 0.003 | *** | |
| D | Intercept | -0.69 | 0.38 | / | 0.44 | / | 0.07 / |
| | $VP_{cur}$ | 3.24 | 0.34 | *** | 0.42 | *** | |
| | $cos_{doy}$ | 0.70 | 0.41 | ** | 0.27 | *** | |
| | $res_{VPD}$ | -2.11 | 0.22 | *** | 0.24 | ** | |

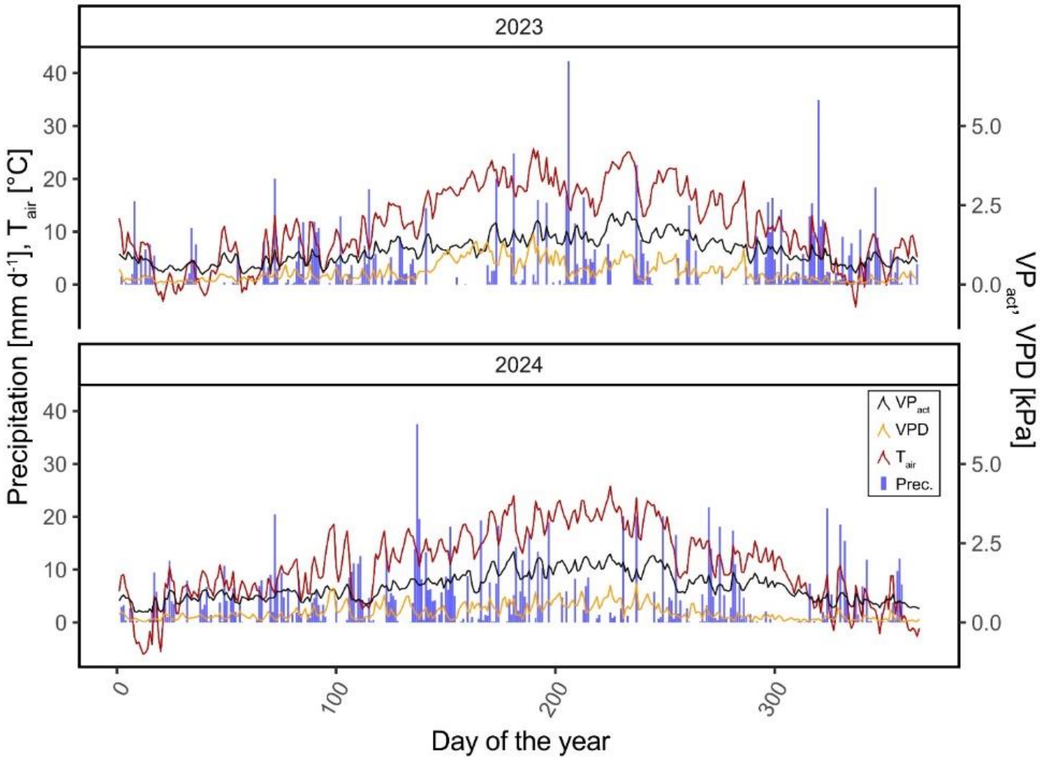


*Figure S2: Meteorological data during the two-year measurement period.*

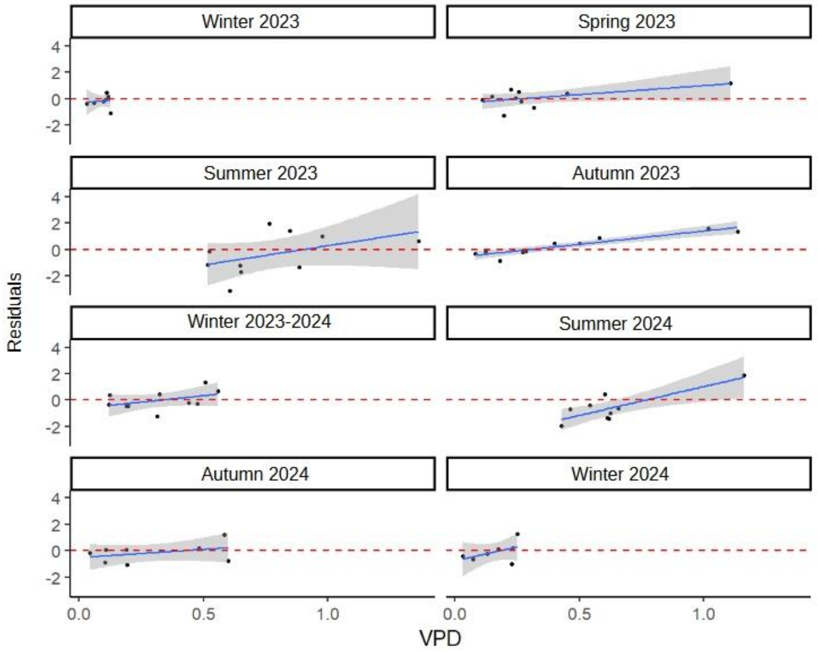


*Figure S3: Residuals of the meteorological model show systematic correlations with the vapour pressure deficit (VPD).*

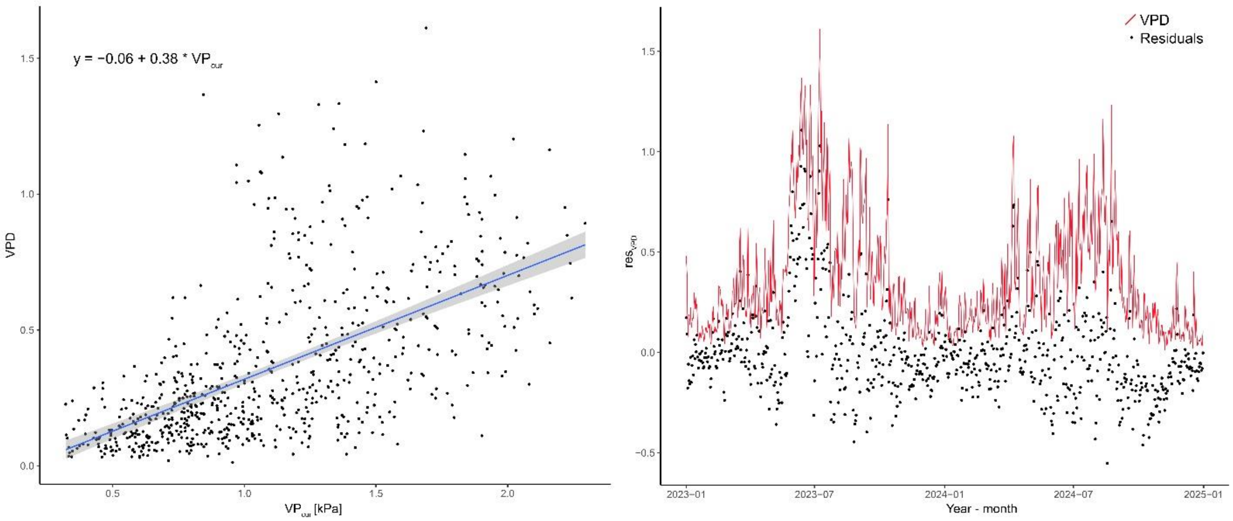


*Figure S4: Linear model explaining the variability in VPD using the VPcur (left) with residuals of the model ($res_{VPD}$) as new input predictor for the meteorological model.*

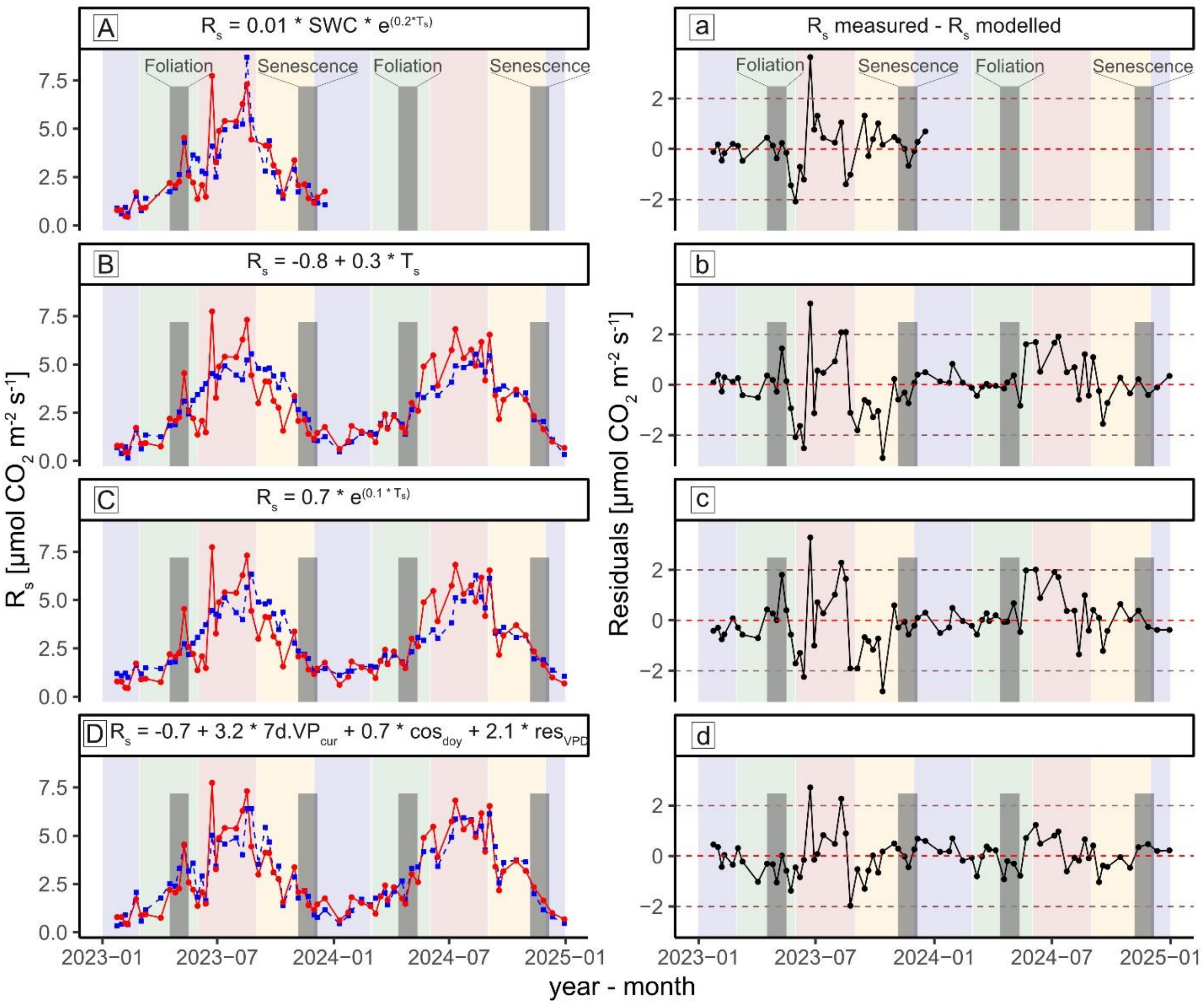


*Figure S5: Figure 3: Temporal model (A-D) for $R_s$ [µmol $CO_2$ $m^{-2}$ $s^{-1}$] for both measurement years in comparison to measured $R_s$ (daily averaged) and absolute residuals of the corresponding model (a-d) for the whole measurement period. Colored areas represent seasons (Blue = winter, green = spring, red = summer, orange = autumn).*

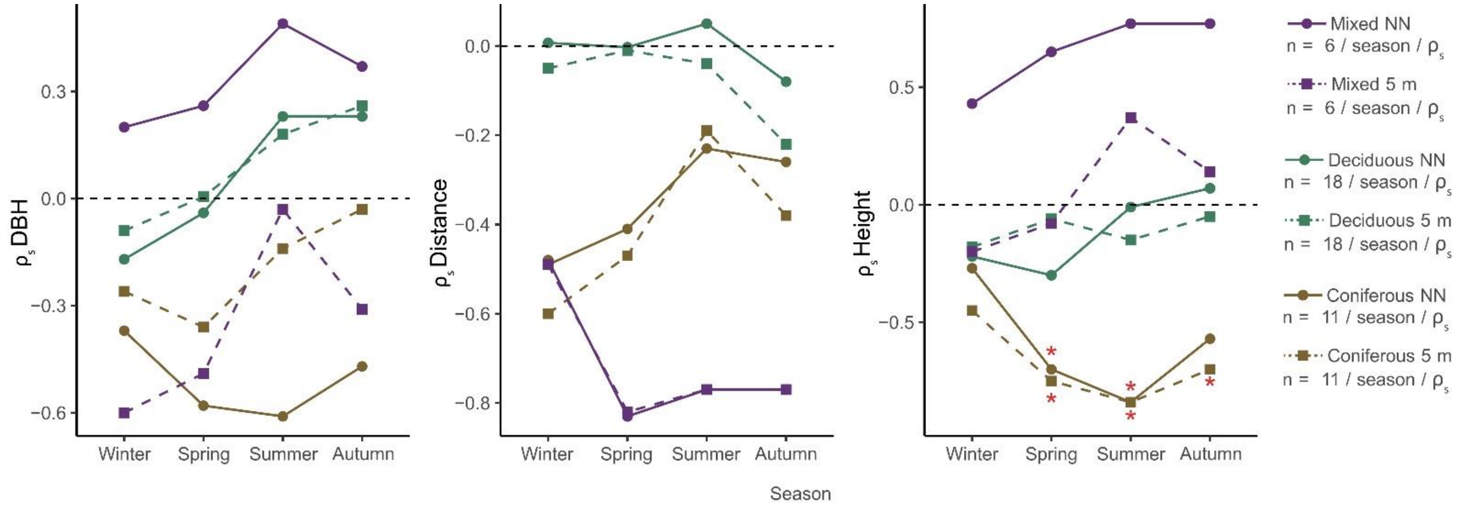


*Figure S6: Spearman correlation coefficients between soil respiration and forest stock features. Soil respiration was averaged per season over the 2-year investigation period. Forest stock features are presented for the nearest neighbouring tree (NN) and the average of the feature of all trees within 5 m buffer zone around measurement station. Significant coefficients are highlighted with * ($p < 0.05$).*

### Publication bibliography


Epron, Daniel; Farque, Lætitia; Lucot, Éric; Badot, Pierre-Marie (1999): Soil CO2 efflux in a beech forest: dependence on soil temperature and soil water content. In *Ann. For. Sci.* 56 (3), pp. 221–226. DOI: 10.1051/forest:19990304.

Gutachterausschuss Forstliche Analytik (2022): Handbuch Forstliche Analytik: Eine Loseblatt-Sammlung der Analysemethoden im Forstbereich. Bundesministerium für Verbraucherschutz, Ernährung und Landwirtschaft, Bonn. Base edition, June 2005, with supplements, 6th edition (February 2022). Available online at https://blumwald.thuenen.de/fileadmin/blumwald/BZE/HFA_Gesamtdatei_2022.pdf, checked on 6/17/2025.

Hartge, Karl Heinrich; Horn, Rainer (1989): Die physikalische Untersuchung von Böden. Stuttgart, Germany: Enke (Vol. 2). Available online at https://library.wur.nl/webquery/titel/505652.

Hereș, Ana-Maria; Bragă, Cosmin; Petritan, Any Mary; Petritan, Ion Catalin; Curiel Yuste, Jorge (2021): Spatial variability of soil respiration (R s ) and its controls are subjected to strong seasonality in an even-aged European beech (Fagus sylvatica L.) stand. In *European J Soil Science* 72 (5), pp. 1988–2005. DOI: 10.1111/ejss.13116.

Knohl, Alexander; Søe, Astrid R. B.; Kutsch, Werner L.; Göckede, Mathias; Buchmann, Nina (2008): Representative estimates of soil and ecosystem respiration in an old beech forest. In *Plant and Soil* 302 (1-2), pp. 189–202. DOI: 10.1007/s11104-007-9467-2.

Kühne, Anke; Schack-Kirchner, Helmer; Hildebrand, Ernst E. (2012): Gas diffusivity in soils compared to ideal isotropic porous media. In *Journal of Plant Nutrition and Soil Science* 175 (1), pp. 34–45. DOI: 10.1002/jpln.201000438.

Ruehr, Nadine K.; Buchmann, Nina (2010): Soil respiration fluxes in a temperate mixed forest: seasonality and temperature sensitivities differ among microbial and root-rhizosphere respiration. In *Tree physiology* 30 (2), pp. 165–176. DOI: 10.1093/treephys/tpp106.

Ruehr, Nadine K.; Knohl, Alexander; Buchmann, Nina (2010): Environmental variables controlling soil respiration on diurnal, seasonal and annual time-scales in a mixed mountain forest in Switzerland. In *Biogeochemistry* 98 (1-3), pp. 153–170. DOI: 10.1007/s10533-009-9383-z.

Saiz, Gustavo; Green, Carly; Butterbach-Bahl, Klaus; Kiese, Ralf; Avitabile, Valerio; Farrell, Edward P. (2006): Seasonal and spatial variability of soil respiration in four Sitka spruce stands. In *Plant and Soil* 287 (1-2), pp. 161–176. DOI: 10.1007/s11104-006-9052-0.

Schack-Kirchner, Helmer; Gaertig, Thorsten; v. Wilpert, Klaus; Hildebrand, Ernst E. (2001): A modified McIntyre and Phillip approach to measure top-soil gas diffusivity in-situ. In *Journal of Plant Nutrition and Soil Science* 164 (3), pp. 253–258. DOI: 10.1002/1522-2624(200106)164:3<253::AID-JPLN253>3.0.CO;2-G.

Søe, Astrid R. B.; Buchmann, Nina (2005): Spatial and temporal variations in soil respiration in relation to stand structure and soil parameters in an unmanaged beech forest. In *Tree physiology* 25 (11), pp. 1427–1436. DOI: 10.1093/treephys/25.11.1427.

Trüby, Peter; Aldinger, Eberhard (1989): Eine Methode zur Bestimmung austauschbarer Kationen in Waldböden. In *J Plant Nutrition & Soil* 152 (3), pp. 301–306. DOI: 10.1002/jpln.19891520307.
